# A Bayesian semi-parametric model for thermal proteome profiling

**DOI:** 10.1101/2020.11.14.382747

**Authors:** Siqi Fang, Paul D.W. Kirk, Marcus Bantscheff, Kathryn S. Lilley, Oliver M. Crook

## Abstract

The thermal stability of proteins can be altered when they interact with small molecules, other biomolecules or are subject to post-translation modifications. Thus monitoring the thermal stability of proteins under various cellular perturbations can provide insights into protein function, as well as potentially determine drug targets and off-targets. Thermal proteome profiling is a highly multiplexed mass-spectrommetry method for monitoring the melting behaviour of thousands of proteins in a single experiment. In essence, thermal proteome profiling assumes that proteins denature upon heating and hence become insoluble. Thus, by tracking the relative solubility of proteins at sequentially increasing temperatures, one can report on the thermal stability of a protein. Standard thermodynamics predicts a sigmoidal relationship between temperature and relative solubility and this is the basis of current robust statistical procedures. However, current methods do not model deviations from this behaviour and they do not quantify uncertainty in the melting profiles. To overcome these challenges, we propose the application of Bayesian functional data analysis tools which allow complex temperature-solubility behaviours. Our methods have improved sensitivity over the state-of-the art, identify new drug-protein associations and have less restrictive assumptions than current approaches. Our methods allows for comprehensive analysis of proteins that deviate from the predicted sigmoid behaviour and we uncover potentially biphasic phenomena with a series of published datasets.

## 1 Introduction

Thermal proteome profiling (TPP (Savitski *et al*., 2014), also referred to as MS-CETSA) is a multiplexed mass-spectrometry extension of the cellular thermal shift assay (CETSA (Molina *et al*., 2013; Jafari *et al*., 2014)). The guiding principle of these experiments is that heating generally causes proteins to denature and become insoluble. This heating can be performed at various temperatures and the remaining soluble protein quantified by mass-spectrometry (MS). This allows a temperature-solubility relationship to be determined and this is frequently called a melting curve (Savitski *et al*., 2014). The melting curve for each proteins is context specific and can be modulated upon binding to small molecules (Gad *et al*., 2014; Huber *et al*., 2015; Chan-Penebre *et al*., 2015). Thus by determining this melting curve for a large number of proteins in different contexts, for example in the presence of a drug, one can find targets and off targets of these molecules (Savitski *et al*., 2014).

There are numerous applications of TPP and it is most commonly used to decipher drug-protein behaviours (Savitski *et al*., 2014, 2018; Huber *et al*., 2015; Reinhard *et al*., 2015; Becher *et al*., 2016, 2018; Mateus *et al*., 2016, 2018). Moreover it can be applied to study interactions with metabolites, nucleotides and nucleic acids (Saei *et al*., 2018; Dziekan *et al*., 2019; Sridharan *et al*., 2019; Becher *et al*., 2018). Authors have shown that proteins in complex with each other are more likely to have concordant *in vivo* melting curves (Tan *et al*., 2018) and others have demonstrated that phosphorylation can alter thermal stability (Huang *et al*., 2019; Potel *et al*., 2020; Smith *et al*., 2020). Thermal proteome profiling has also been complemented with extensive structural analysis (Feng *et al*., 2014; Leuenberger *et al*., 2017; Schopper *et al*., 2017; Piazza *et al*., 2018). Furthermore, TPP is not just applicable in human cells but can be applied in bacteria *in vivo* (Mateus *et al*., 2018), in the apicomplexan parasite *Plasmodium falciparum* (Dziekan *et al*., 2019, 2020), and in tissue or blood (Perrin *et al*., 2020). Extensive work has recently been presented characterising the melting behaviour of proteins across 13 species, demonstrating similarities and difference for protein orthologues (Jarzab *et al*., 2020).

Thermodynamic theory predicts that the melting curve of proteins should have a sigmoid behaviour (Schellman, 1994). Melting curves of a protein may then be compared to determine context specific behaviours. Statistical analysis can then follow a number of directions. For example, one approach involves summarising melting curves into a *T_m_* - the temperature at which relative solubility has halved (Savitski *et al*., 2014; Huber *et al*., 2015). This is then followed by comparison of *T_m_* values across the two contexts using the appropriate *z*-score. This approach assumes that the melting curve is a bijection, else there might be multiple candidates for *T_m_*. It also assumes that *T_m_* is defined, which need not be the case if relative solubility has never halved. Another approach is to compare the relative solubility at a fixed temperature (Ball *et al*., 2020). However, summarising curves to a single value results in loss of information, loss of sensitivity and does not account for the quality of the fit of the parametric model (Childs *et al*., 2019). A more powerful approach is to employ techniques from functional data analysis (Ramsay and Dalzell, 1991; Ramsay, 2004; Wang *et al*., 2016) and use the whole melting curve for statistics (Childs *et al*., 2019).

Childs *et al*. (2019) introduced the method *non-parametric analysis of response curves* (NPARC) for powerful analysis of melting curves. In brief, the method assumes a sigmoid model for the data and then proceeds to perform an analysis of variance (ANOVA). Since typically TPP data involves measurement of melting curves for a great many proteins per experiment, the appropriate null distribution can be directly estimated from the data (Efron, 2004, 2012). NPARC allowed thousands more proteins to be analysed that the original *T_m_* centric analysis and demonstrated a significant improvement in statistical power. However, this method still assumes a parametric sigmoid model and the method used to estimate the null distribution assumes that it is unimodal. Moreover, large-scale testing frameworks assume that the large majority of observations are samples from the null distribution, which can be problematic if the context of interest affects many proteins. Furthermore, there is no uncertainty quantification in the melting curves or the key model parameters.

To overcome these limitations, here we develop a Bayesian version of the sigmoid model, which allows uncertainty quantification. Furthermore, in the Bayesian framework one does not need to estimate the null distribution and multiplicity control is automatic via the prior model probabilities (Scott and Berger, 2006, 2010; Berger *et al*., 2014; Chang *et al*., 2020). In addition, including prior information on the model parameters has a number of benefits; such as, allowing the shrinkage of residuals towards 0, the regularisation of the inferred parameters and improved algorithmic stability (Gelman *et al*., 2013). Through exploratory data analysis and model criticism, we find evidence for model expansion. We show that the standard sigmoid model is insufficient to model the relationship between temperature and relative solubility for some proteins. This motivates the development a semi-parametric model (Powell, 1994). A semi-parametric model is one that includes both parametric terms, in our case the sigmoid, and unknown non-parametric terms. A *Gaussian Process prior* (GP prior, (Stein, 2012)) is used to infer the non-parametric terms. Gaussian processes are highly flexible and have been used extensively in other molecular biology applications, such as gene-expression time courses (Kirk and Stumpf, 2009; Kirk *et al*., 2012; Stegle *et al*., 2010; Cooke *et al*., 2011; Babtie *et al*., 2014), single-cell transcriptomics (Reid and Wernisch, 2016; Boukouvalas *et al*., 2018; Strauss *et al*., 2020) and spatial proteomics (Crook *et al*., 2019; Shin *et al*., 2019).

Here we begin with exploratory data analysis of five datasets which motivates the creation of more flexible models. We then carefully analyse published data to demonstrate the improved sensitivity our method, as well as the value of uncertainty quantification. We identify putative protein-drug interactions that have been overlooked in previous TPP studies, including the protein HDAC 7 in studies designed to determine targets of the chemotherapeutic drug, Panobinostat. We proceed to characterise the proteins that deviate from sigmoid behaviour and uncover functional, as well as localisation, enrichments.

## 2 Results

### 2.1 Exploratory data analysis motivates model extension

First, we interrogated data from five TPP experiments that were performed on the K562 human erythroleukemia cell line. The first experiment explored the effects of detergents on ATP-binding profiles. Then two other experiments explored the effects of different concentrations of the ABL inhibitor Dasatinib. In one of the experiments the histone deacetylase (HDAC) inhibitor Panobinostat was used to determine its effects on the behaviour of proteins. The final experiment explored the effects of the pan-kinase inhibitor Staurosporine. A summary of the experiments is given in table 1.

**Table 1:** Summary of the datasets and the respective reference used in this manuscript.

| Dataset | Treatment | Concentration | number of proteins | Reference | Intact or Lysate |
| --- | --- | --- | --- | --- | --- |
| ATP data | MgATP | 2 $\mu M$ | 4177 | Reinhard *et al.* (2015) | Lysate |
| Dasatinib 0.5 data | Dasatinib | 0.5 $\mu M$ | 4625 | Savitski *et al.* (2014) | Intact |
| Dasatinib 5 data | Dasatinib | 5 $\mu M$ | 4154 | Savitski *et al.* (2014) | Intact |
| Panobinostat data | Panobinostat | 1 $\mu M$ | 3649 | Franken *et al.* (2015) | Intact |
| Staurosporine data | Staurosporine | 20 $\mu M$ | 4505 | Savitski *et al.* (2014) | Lysate |

We applied the NPARC pipeline to each of these experiments and carefully explored the results. The NPARC analysis approach makes a number of assumptions. Firstly, when estimating the null distribution, it assumes that the it is unimodal and thus a single F distribution is appropriate to approximate the null distribution. Secondly, it assumes that a large majority of the observed data are samples from the null distribution, which might not be the case for some contexts. Finally, it assumes that the sigmoid model is appropriate. To clarify, the 3-parameter sigmoid model of interest is the following:

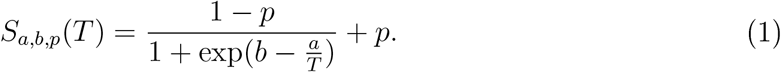

The parameter *p* is interpreted as a plateau, whilst *a* and *b* are shape parameters. This sigmoid model, and more generally sigmoid functions, makes the assumption of monotonicity, a single inflexion point, rotational symmetry around the inflexion point, a bell-shaped first derivative and horizontal asymptotes (at *p* and 1 − *p*). In many cases, such assumptions are appropriate and this behaviour is widespread in the TPP datasets we examined (see figure 1 C and E). However, we did observe proteins which deviated from this behaviour and violated these assumptions (between 3 and 20% depending on the dataset), beyond what could be attributed to measurement error. These include examples of a hyper-solubilisation phenomena; that is, proteins reproducibly increasing in relative solubility as temperature increases, which is not predicted by thermodynamics (Schellman, 1994).

**Figure 1:**
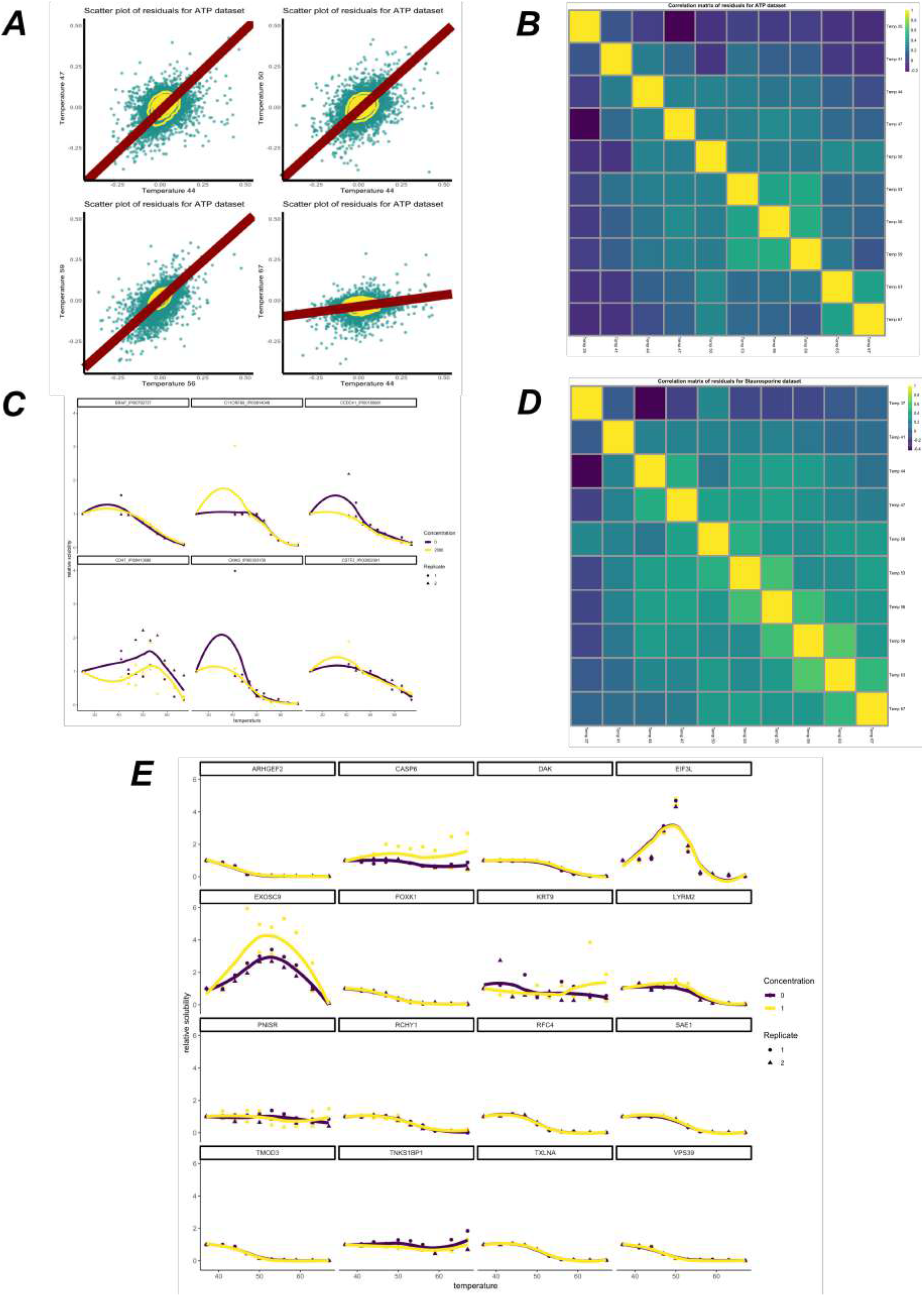
(A) Scatter plots of residuals for the sigmoid model at different temperatures applied to the ATP dataset (Reinhard *et al*., 2015). Orthogonal regression line shown in dark red and contours shown in yellow. Residuals are strongly correlated. (B) Sample Spearman correlation matrix of the the residuals for the ATP dataset. (C) Example melting curves for some proteins from the ATP dataset. LOESS curves shown for visualisation. (D) as for B, but for the Staurosporine dataset (Savitski *et al*., 2014). (E) Example melting curves from the Panobinostat dataset (Franken *et al*., 2015). LOESS curves shown for visualisation.

After fitting a sigmoid model to each protein in each condition, we computed the residuals for every protein at each temperature. Classical analysis of variance assumes that the residuals are independently and normally distributed with homoscedasticity. We observed that none of these conditions are true for these data (see figure 1 A for an example). Childs *et al*. (2019) also noted this fact by comparing the empirically derived F distributions to those which would be obtained under classical assumptions and by also analysing the corresponding *p*-value histograms (Holmes and Huber, 2018). Significant departure of the F distributions from the theoretical behaviour was observed and so they used large scale data analysis tools to approximate the null. This results in different effective degrees of freedom for the F test and analysis of variance proceeds as usual. For sake of pedagogy, we state that bootstrapping or permutation methods, amongst others, could also have been used (Efron, 2012).

To perform residual analysis, we computed the sample Spearman correlation matrix for the residuals and observed that different datasets have different correlation structures (see figures 1 B and C) and that residuals for closer temperatures are, in general, more correlated. The presence of correlated residuals usually suggests data structure that has not been correctly modelled (Glasbey, 1979, 1980; Crowder and Hand, 1990).

To avoid estimating the null distribution, we recast the analysis of TPP data by proposing a Bayesian sigmoid model. This has the further benefit of allowing expert prior information to be included for the parameters. The Bayesian framework also allows us to quantifying the uncertainty in our parameter estimates and as a result the uncertainty in the fitted function. Given that we observed deviations from the sigmoid model and strongly correlated residuals, we proposed to include an additional functional term in our model. Given no suitable parametric candidate for this additional term, we sought inspiration from the Bayesian non-parametric literature and placed a *Gaussian process prior* on this additional term, allowing a more flexible set of functions to be modelled and the uncertainty in this function to be quantified (Dudley *et al*., 1973; Rasmussen, 2003; Ghosh and Ramamoorthi, 2003). We refer to the methods section for a precise description of our model.

### 2.2 Analysis of Staurosporine dataset

Having developed sigmoid and semi-parametric Bayesian models, we applied these approaches to the Staurosporine dataset (Savitski *et al*., 2014). Staurosporine is a pan-kinase inhibitor, where the inhibition is achieved by a having high affinity to the ATP-binding site of kinases (Karaman *et al*., 2008). How Staurosporine affects the cell is not completely understood and has been shown to induce apoptosis (Chae *et al*., 2000) and cell cycle arrest (Bruno *et al*., 1992). The Staurosporine dataset that we consider reports relative solubility of proteins in the presence of 20*μM* of Staurosporine for 2 control replicates and 2 treatment replicates. A total of 4505 proteins were measured using quantitative multiplexed TMT LC-MS/MS measurements at temperatures ranging from 37 degrees to 67 degrees in 10 even increments of 3 degrees (Savitski *et al*., 2014).

One advantage of this dataset is that we expect a large number of kinases to be the target of Staurosporine. Hence, we might expect such proteins to have shifts in their thermal profiles upon Staurosporine treatment. Hence, as in previous analysis (Childs *et al*., 2019), we curate a set of proteins with the annotation “protein kinase activity” from *ensembl.db* (Zerbino *et al*., 2018). We then compute the sensitivity, the proportion of correctly identified positive cases, for the NPARC and two Bayesian, sigmoid and semi-parametric, approaches (taking the *p*-value threshold as 0.01 and, equivalently, the posterior probability threshold as 0.99). The NPARC approach achieves a sensitivity of 33.7, whilst the Bayesian sigmoid model a sensitivity of 36.7 and the Bayesian semi-parametric model achieves 39.6 (see figure 2 B). This suggests that avoiding estimation of the null and expanding the model flexibility can improve the sensitivity of the analysis. Unfortunately, in such cases specificity (the true negative rate), is not well defined, since proteins that are not kinases may also have their melting curve perturbed, perhaps due to changes in their phosphorylation state as a result of ablated kinase function (Potel *et al*., 2020). We see similar improvements for sensitivity when considering other datasets (see supplement) and a simulation study is also included in the supplement demonstrating that the two Bayesian approaches outperform the NPARC method.

**Figure 2:**
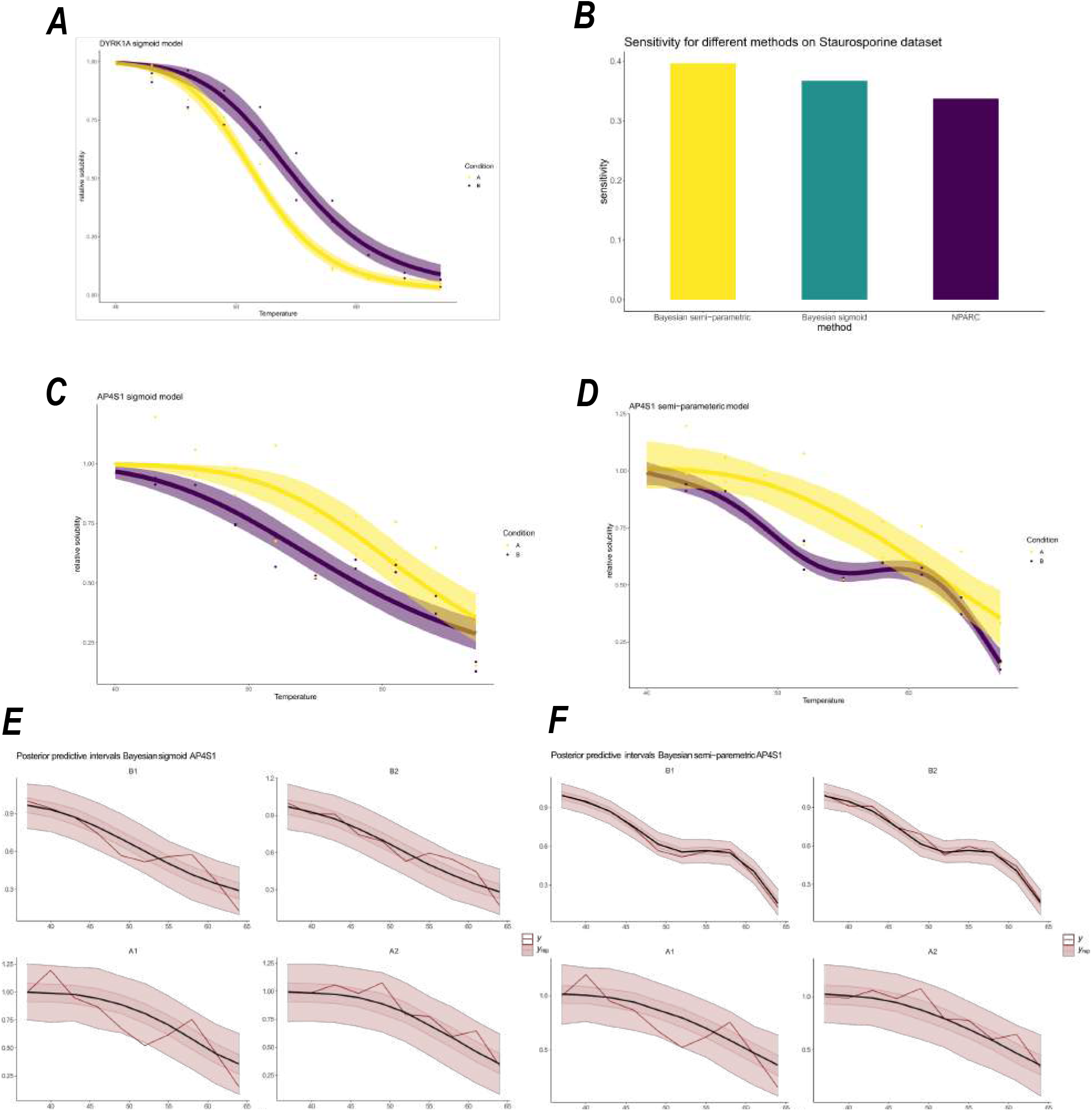
(A) Melting profile for the DYRK1A with inferred mean sigmoid model function plotted, along with 95% credible bands for the inferred mean function. (B) Sensitivity for the different methods applied to the Staurosporine dataset (c) Melting profile for AP4S1 using the sigmoid model, with uncertainty estimates in mean function (D) Melting profile for AP4S1 using the semi-parametric model, including inferred mean function and 95% confidence bands. (E,F) Posterior predictive checks for AP4S1 using the two Bayesian models: (E) sigmoid (F) semi-parametric. The red line correspond to the observed data. Whilst the black line is the posterior predictive mean function and the credible bands correspond to 50% and 95% credible bands of the posterior predictive distribution, respectively.

Improved sensitivity results in finding new proteins that are putative targets of Staurosporine. For example, DYRK1A, a dual-specificity kinase with both serine and tyrosine kinase activities (Shindoh *et al*., 1996; Papadopoulos *et al*., 2011), which is essential for brain development (Ogawa *et al*., 2010; Soundararajan *et al*., 2013), was overlooked by the NPARC analysis. Our Bayesian analysis is able to determine DYRK1A as a kinase which is stabilised by Staurosporine (posterior probability > 0.99). This observation is supported by kinobeads competition-biding experiments, where DYRK1A demonstrated a Staurosporine dependent effect (pIC_50_ = 6.58) (Werner *et al*., 2012) and an isothermal shift assay (iTSA) also demonstrated a Staurosporine dependent effect on DYRK1A at 52°C (Ball *et al*., 2020). Figure 2 A demonstrates other benefits of the Bayesian analysis, where we visualise uncertainty in the inferred sigmoid mean function. There is clear separation between the sigmoid curve between the two conditions. However, it also highlights the potential limitations of the sigmoid model, with rotational symmetry imposed around the point of inflexion.

An even clearer example were the sigmoid model fails is the case of AP4S1, a component of the adaptor protein complex which is involved in vesicle trafficking from the trans-Golgi to the endosome (Hirst *et al*., 1999; Del?Angelica *et al*., 1999). Figure 2 C shows the sigmoid model cannot model the multiple inflexion points of the melting curve of AP4S1. The limitation being the single inflexion point. Figure 2 D shows the inferred mean function and associated uncertainty estimates. Clearly the semi-parametric model is more appropriate for such cases.

To compare these models more formally, we performed a *posterior predictive check* (see section 4.2). From the posterior predictive distributions, we examined the credible bands. To be precise, given a model, an observed value is predicted to fall in the credible band of size *β* with probability *β*. Hence, if the observed data fall outside the credible bands, it is indicative of the model being insufficient. From figure 2 E we see the data frequently lies outside the 50% credible band and occasionally outside the 95% credible band. Whilst for the semi-parametric model, visualised in figure 2 F, the data never falls outside the 95% credible band and is more frequently contained in the 50% credible band. This suggests that the semi-parametric is more appropriate, in the this case. Kernel density estimate based posterior predictive checks make a similar conclusion and are included in the supplement.

For a more quantitative treatment, we examine the out-of-sample predictive accuracy from the fitted Bayesian models (see section 4.2). We use leave-one-out cross validation (LOO-CV) with the log-predictive density as the utility function. Higher scores indicate better out-of-sample predictive performance. The LOO-CV estimate for the sigmoid model is 26.7 ± 5.4(SE), whilst for the semi-parametric model it is 41.1 ± 6.5(SE). We conclude, for this protein (AP4S1), the semi-parametric model is superior. As a result of the improved modelling, our analysis was able to determine that AP4S1 was destabilised upon Staurosporine treatment (posterior probability > 0.99), which we could not determine from NPARC or the Bayesian sigmoid model. AP4S1 is not a kinase, thus its change in behaviour upon Staurosporine treatment is not straightforward to interpret. In any case, we would expect kinases to be stabilised, rather than destabilised. This destabilisation might be an effect of not being correctly localised or not being able to correctly form a complex. AP4S1 localisation is dependent on the small G protein ARF1 (Yu *et al*., 2012), whose function, it turn, depends on several kinases (Rümenapp *et al*., 1997; Morohashi *et al*., 2010). Thus, the destabilisation is likely a downstream effect of Staurosporine as a pan-kinase inhibitor.

### 2.3 Proteins with altered thermal stability upon Panobinostat treatment

The analysis of the Staurosporine dataset demonstrated the improved sensitivity of our method and the ability of our approaches to model complex behaviours, whilst also quantifying uncertainty. We next applied our method to the Panobinostat dataset where, in the original analysis, only a handful of hits were identified (Franken *et al*., 2015). Panobinostat is a non-selective histone deacetylase inhibitor (pan-HDAC inhibitor) that is approved for use in patients with multiple myeloma (Laubach *et al*., 2015). Thermal proteome profiling was applied to K562 cells treated with a vehicle (control) or 1*μM* of Panobinostat. 2 replicates in each context were produced and a total of 3649 proteins were measured (Franken *et al*., 2015). These panobinostat experiments are cell-based rather than lysates and so we expect our approach to be sensitive to non-canoncial melting curves that may be due to effects on solubility.

We applied the NPARC pipeline and identified 7 proteins as having their melting curve significantly altered (*p* < 0.01), which included the known Panobinostat targets HDAC 1, 6,8,10. The HDAC proteins are responsible for deacetylation of lysine residuse of the N-terminal of the core histones, as well as other proteins (Grozinger *et al*., 1999; Hubbert *et al*., 2002; Seto and Yoshida, 2014; Allis and Jenuwein, 2016; Li and Seto, 2016). To quantify uncertainty, we applied the Bayesian sigmoid approach, also avoiding estimation of the null distribution. The Bayesian sigmoid model was able to identify 34 proteins whose melting profile was treatment dependent (posterior probability > 0.99). 16 of these proteins are plotted in figure 3 and these putative hits included all of the proteins discovered by the NPARC approach.

**Figure 3:**
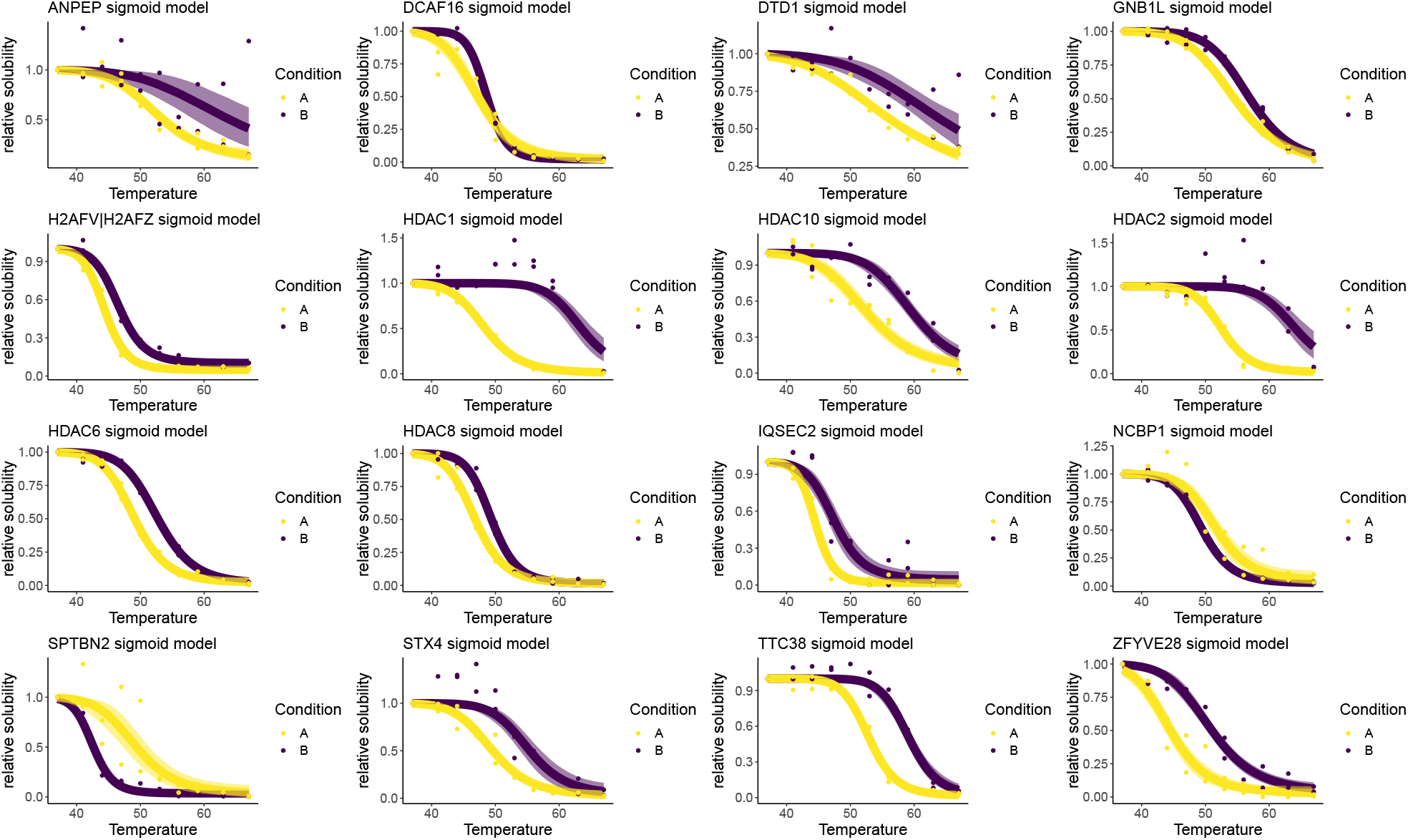
Melting profiles for 16 protein with posterior probability > 0.99 in favour of a condition dependent model using the Bayesian sigmoid model. Points are observed protein measurements. The inferred mean function from the sigmoid model is plotted as a line and the 95% credible band is given by the shaded region. Purple denotes the drug treated context, whilst yellow denotes the control.

We also observed several proteins whose melting behaviour was not previously known to depend on Panobinostat; such as, NCBP1 whose behaviour appears to be destabilised upon Panobinostat treatment. NCBP1 is a nuclear cap-binding protein that is dual localised to the cytosol and nucleus, as well as being an integral component of the cap-binding complex (Izaurralde *et al*., 1994, 1995). Given the role of acetylation in formation of protein complexes (Choudhary *et al*., 2009), as well as NCBP1 having been shown to have two lysine residues that are substrates for acetylation (Choudhary *et al*., 2009) it possible that the observed melting behaviour is a downstream result of the ablated function of the HDAC proteins.

We have already demonstrated that non-sigmoidal behaviour is not unusual in the Panobinostat dataset (see figure 1 E). Hence, we applied our Bayesian semi-parametric model to these data. We identified 85 proteins whose melting profile was panobinostat dependent with posterior probability greater than 0.99. These included HDAC 7, one of the core members of the histone deacetylation complex, which was not identified by either NPARC or the Bayesian sigmoid model (Figure 4). In this case, however, HDAC 7 is not stabilised but, rather, destabilised suggesting indirect regulation downstream of Panobinostat targets. This finding is consistent with a recent report showing that HDAC 7 abundance is regulated through activity of the known Panobinostat targets HDAC 1 and 3 (Caslini *et al*., 2019) and with HDAC 7 not being enriched in pull-down experiments with the Panobinostat (Becher *et al*., 2016).

**Figure 4:**
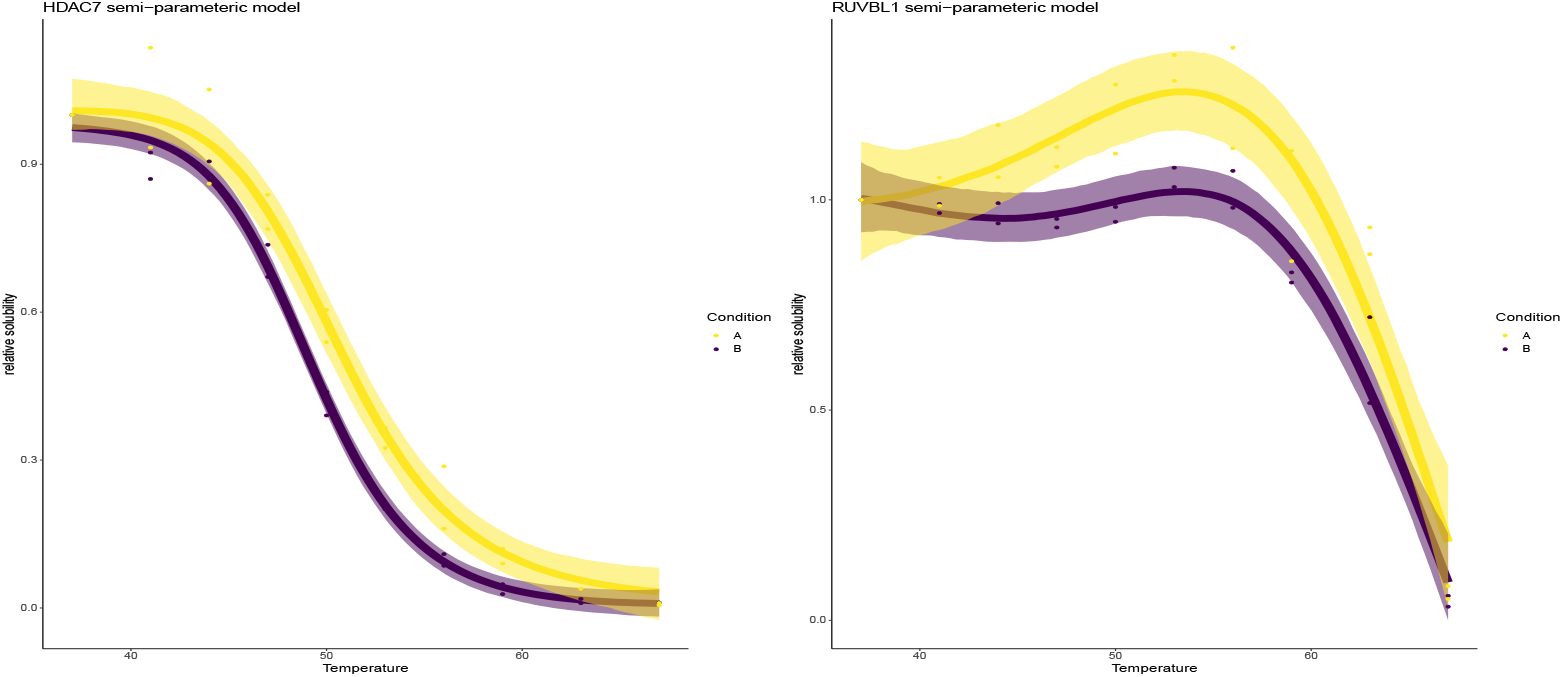
Melting profiles for HDAC 7 and RUVBL1 using the Bayesian semi-parametric model. The points are observed protein data. The line represents the inferred mean function and the shaded region is the 95% credible band for the inferred mean function. Purple denotes the drug treated context, whilst yellow denotes the control.

Another protein which we identified with Panobinostat dependent behaviour was RU-VBL1. RUVBL1 is a well studied protein involved in histone acetylation and is a component of several complexes, has multiple localisations and many interaction partners (Doyon *et al*., 2004; Cai *et al*., 2003; Wood *et al*., 2000; Ikura *et al*., 2000; Cloutier *et al*., 2017; Obri *et al*., 2014). RUVBL1 displays curious behaviour with both hypersolubilsation and destabilisation upon treatment with Panobinostat (Figure 4). Since RUVBL1 has multiple states and is involved in multiple different complexes, it is possible that the effects of Panobinostat are interrupting only a certain pool of RUVBL1 proteins, leading to biphasic behaviour. Certain functional units of RUVBL1 might be more thermally stable than others, leading to complex temperature-solubility behaviours. The extent to which the behaviours are reflected in the melting curves will depend on many factors. Two dimensional thermal profiling experiments in lysate HepG2 cells show that RUVBL1 is highly thermal stable and did not display sigmoidal behaviour at several concentrations of Panobinostat (5,1, 0.143, 0.02)*μM* at a temperature range of 42 – 63.9°C (Becher *et al*., 2016).

### 2.4 Characterising proteins that deviate from sigmoid behaviour

Having established the utility of our Bayesian models, in particular the ability of our semi-parametric approach to model deviations from sigmoid behaviour. We next considered those proteins that were better modelled by the semi-parametric approach to see if they have any physical, functional or otherwise defining features. We began our investigation by selecting a set of proteins where the semi-parametric model explains at least 5% more variance (Gelman *et al*., 2019) than the sigmoid model does alone.

We performed functional enrichment testing of these proteins using UniprotKB annotations. We found that the posttranslation modifications acetlyation and phosphoprotein are enriched in these proteins across the 5 human datasets (∀_*i*_, *p_i_* < 10^−8^ Fisher exact BH corrected), as well as RNA binding (∀_*i*_, *p_i_* < 10^−6^ Fisher exact BH corrected). The pattern of enrichment can be visualised in figure 5 A and is reproducible across all the datasets. Whilst the effect of phosphorylation on protein thermal stability is well appreciated (Potel *et al*., 2020), the role of acetylation on thermal stability has not been characterised, despite well established influence on protein stability (Choudhary *et al*., 2009). Enrichment of acetylated proteins could suggest a mechanistic effect of acetylation on thermal stability.

**Figure 5:**
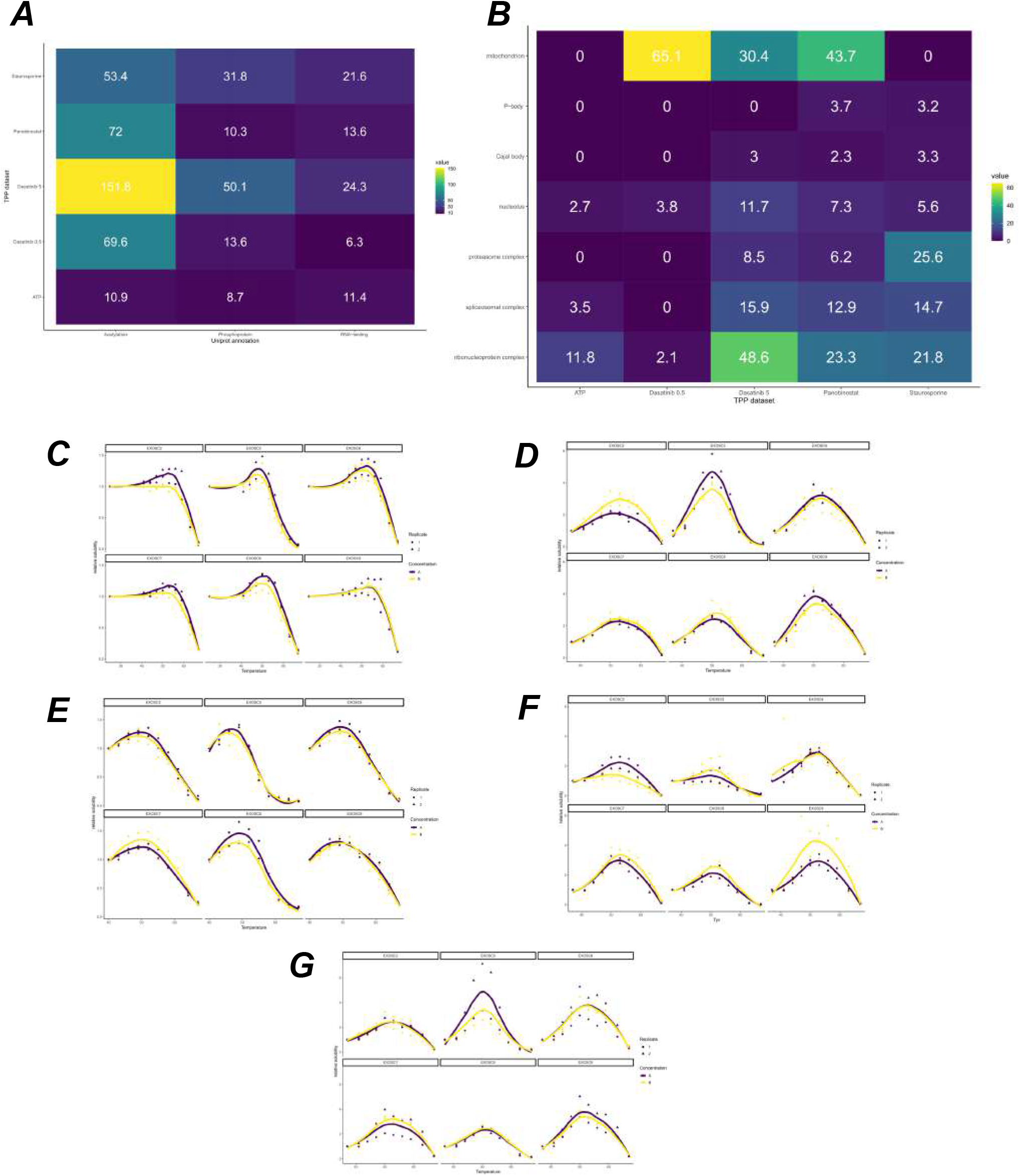
(A) Uniprot key term enrichment analysis. A tile plot show −log_10_ of the *p*-values for each of the terms for the 5 human datasets. (B) GO CC enrichment analysis. A tile plot showing −log_10_ of the *p*-values for each of the terms for the 5 human datasets. (C-G) Melting profiles of the proteins from the EXOSC complex, across the 5 human datasets, (C) ATP dataset (D) Dasatinib 5 dataset (E) Staurosporine dataset (F) Panobinostat dataset (G) Dasatinib 0.5 dataset

Non canonical melting behaviour may represent different pools of the same protein behaving differently within the cell. Non-canonical proteins are enriched for RNA-binding proteins and so the different species of protein, i.e the RNA-bound form or the entities not bound to RNA, might have different temperature-solubility relationships, as well as different drug induced behaviours. Hence, what we may be observing in TPP datasets is a mixture of these behaviours being reflected in different ways. The extent to which one observes such behaviours will depend on the relative number of copies of each protein in each state and also on the the particular way the modification effects the thermal stability of the protein. Hence, exactly which protein display this behaviour will be cell line and context specific, and so requires further investigation. This interpretation would explain both the hypersolubilisation and biphasic behaviour we have observed.

We continued to characterise the subcellular localisations of these proteins, with the hypothesis that these protein might come from a single or perhaps multiple localisations. As we see from figure 5 B, the pattern for subcellular localisation is much less consistent than the pattern for functional enrichment and only the nucleolus and the ribonucleoprotein complex are enriched annotations for protein with non-sigmoidal behaviour in all the human datasets.

The nucleolus is a phase-separated sub-nuclear compartment and is the site of ribosome biogenesis (Boisvert *et al*., 2007). Furthermore, during heat stress molecular chaperones accumulate in the nucleolus to protect unassembled ribosomal proteins against aggregation (Frottin *et al*., 2019). This effect is readily seen within 2 hours at 43 degrees. Despite TPP experiments usually only heating for minutes, we hypothesised that functional role of the nucleolus thus guards against the phenomena that TPP is attempting to induce. To test this hypothesis further, we filtered to proteins that are classed as non-sigmoidal and have known nucleolus annotation. We found that several proteins of the exosome complex EXOSC[2,5-9] fall into this class and are measured completely in all experiments. Figures 5 shows the reproducible non-sigmoidal behaviour. Remarkably, all members of this complex show hypersolublisation and increasing stabilisation until roughly 50 degrees. After 50 degrees the proteins destabilised. Without further experiments, we cannot deduce whether this effect is representative of the whole nucleolus or solely these EXOSC proteins. One alluring explanation could be that RNA dissociates from the EXOSC complex at 50 degrees. Furthermore, we de not observe significant co-aggregation of EXOSC protein in (Thermal Proximity CoAggregation) TPCA data (Tan *et al*., 2018). However, TPCA analysis derives curve similarity from an inverse euclidean distance, which may not be a sufficiently sensitive measure of curve similarity in this case.

Continuing our investigation into subcellular localisation, we integrated our analysis with spatial proteomics data from hyperLOPIT experiments (Mulvey *et al*., 2017). We used hyperLOPIT data from U-2 OS cells, providing information on 4883 proteins to 11 sub-cellular compartments (Thul *et al*. (2017); Geladaki *et al*. (2019) and re-analysed in Crook *et al*. (2020) to reveal 14 compartments). We projected the proteins that deviate from sigmoid behaviour onto the PCA coordinate of the hyperLOPIT data (figure 6). In all datasets, we observed enrichment for nuclear, ribosomal and cytosolic regions, in agreement with our GO enrichment analysis. Furthermore, also in support of the GO enrichment results, we saw strong enrichment for mitochondrial annotations in the two Dasatinib datasets and the Panobinostat dataset. To understand the functional relevance of these proteins, we stratified to the proteins that have mitochondrial annotations according to the hyperLOPIT data.

**Figure 6:**
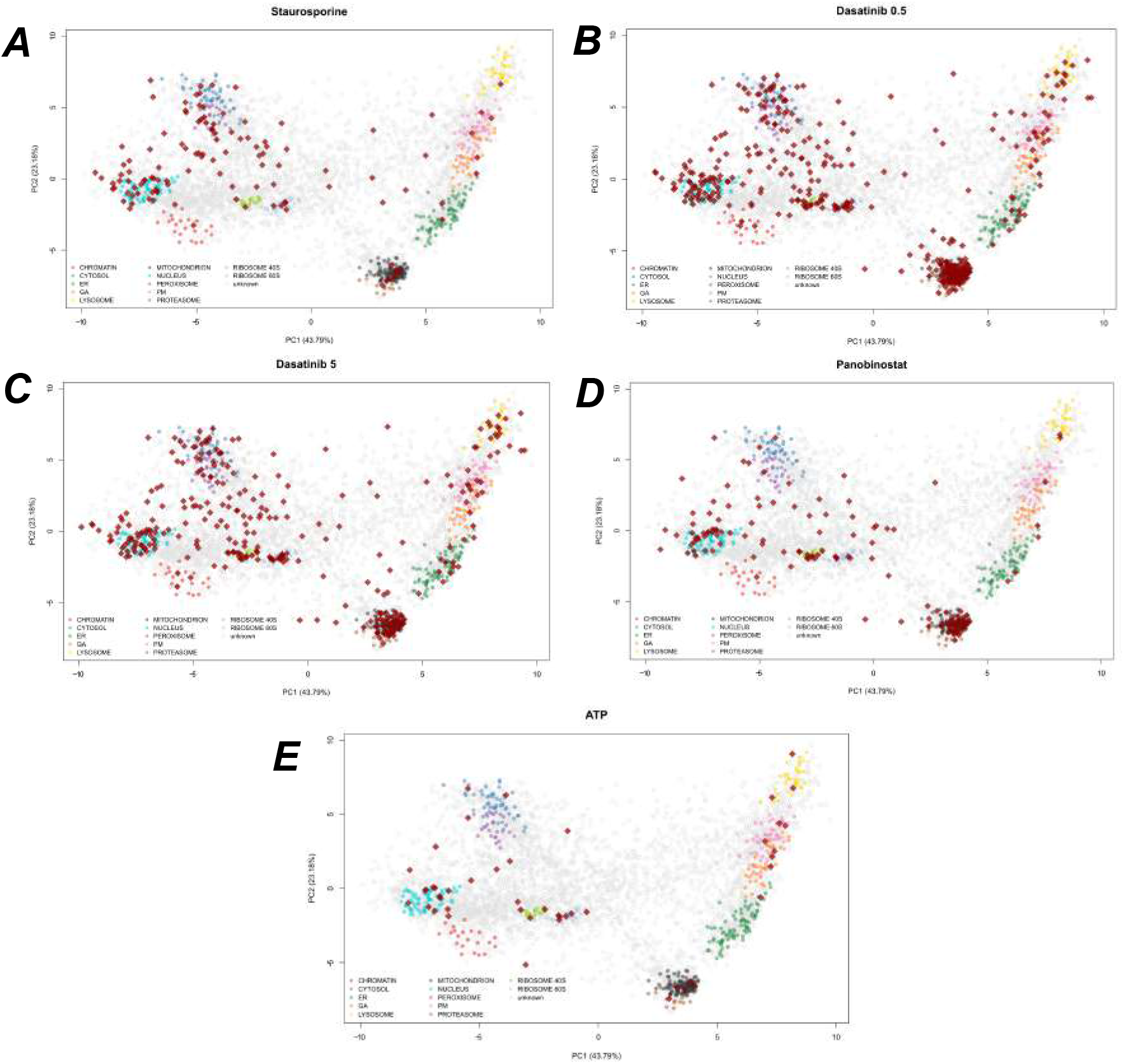
(A-E) PC A plots of U-2 OS hyperLOPIT data (Geladaki *et al*., 2019), showing the top two principal components. Each pointer is a protein and marker proteins for each subcellular niche are coloured. Dark red diamonds denote proteins that were deemed to have non-sigmoid behaviour from TPP data. Each panel represents a different TPP dataset for the projected proteins.

In the Dasatinib 0.5 dataset, we saw enrichment for cofactor binding (*p* < 10^−13^), coenzymee binding (*p* < 10^−9^), NAD binding domains (*p* < 10^−7^), small-molecule binding (*p* < 10^−9^), FAD binding domains (*p* < 0.0001), nucleotide binding (*p* < 10^−9^), ATP-binding and RNA-binding (*p* < 0.05). We see similar results in the Dasatinib 5 dataset: cofactor binding (*p* < 0.001), co-enzyme binding (*p* < 0.001), NAD binding (*p* < 0.001), nucleotide binding (*p* < 0.001), small molecular binding (*p* < 0.01). Almost identical results are seen for the Panobinostat dateset: cofactor binding (*p* < 10^−8^), NAD binding (*p* < 10^−6^), co-enzyme binding (*p* < 10^−6^), small molecule binding (*p* < 0.01), nucleotide binding (*p* < 0.01), FAD binding domain (*p* < 0.01). Taken as a whole, these results support our interpretation of biphasic behaviour where different functional copies of a protein behave differently from each other and that we observe a mixture of these behaviours in TPP experiments.

Given the functional and localisation enrichments we have observed, we sought to further characterise these proteins by examining their intrinsic disorder. Indeed aggregation-prone proteins, after non-lethal heatshock, are enriched for intrinsically disordered regions (Määttä *et al*., 2020). Using the D2P2 database (Oates *et al*., 2012), we first obtained the length of the predicted intrinsically disorder regions (IDRs) for every protein. For stringency, we required that at least a minimum of 4 prediction tools were in agreement. To correct for length bias, we computed the proportion of the protein that was intrinsically disordered. We then tested if the set of proteins with non-canonical melting behaviour were enriched for proteins which had at least 5% of regions predicted to be intrinsically disordered. No such enrichment was observed (Fisher’s exact test). We further filtered to proteins in our analysis that had nucleus annotations and despite nuclear annotated non-canonical proteins having a large proportion of IDRs (80 – 95%), there was no statistical enrichment beyond what one would have expected for nuclear proteins.

A further consideration is whether the experiment was performed in intact or lysed cells. Indeed, for the 3 experiments that were performed on intact cells (Dasatinib 0.5 and 5 and Panobinostat) the non-sigmoidal proteins showed an enrichment for mitochondrial localisation whilst the lysate-based experiments did not. In lysate-based experiments the mitochondrial membrane will break down and the local concentration of NAD will decrease. Hence, the drug has easier access to mitochondrial proteins in lysate-based experiments. Since cellular physiology is preserved for intact cells, we might believe that non-sigmoidal behaviour is a indicative of downstream effects. However, some non-sigmoidal behaviours are reproducible and independent of whether the experiment was in lysed or intact cells. Thus, we cannot completely attribute these effects to whether the experiments were performed in intact cells or not.

## 3 Discussion

We have presented Bayesian approaches to the analysis of thermal proteome profiling data. Our Bayesian sigmoid model quantifies uncertainty and avoids empirical estimation of the null distribution. The resulting model shows improved sensitivity and, as a result, we identified new putative targets and off-targets in 5 human TPP experiments. Uncertainty quantification provides useful additional information and, by inspecting the confidence bands, we can carefully select the temperatures at which to perform validation experiments.

Many proteins exhibit non-sigmoid behaviour and we observed strong correlation between residuals in all the datasets we analysed, motivating an expanded model. Thus, we introduced a semi-parametric Bayesian model that further improved sensitivity, had better out-of-sample predictive properties for some proteins and had confidence bands with improved coverage. This improved analysis allowed us to identify HDAC 7 as having altered thermal stability on Panobinostat treatment, which previous analysis could not identify.

We probed the proteins that deviated from non-sigmoid behaviour and our analysis suggests that these proteins are enriched for proteins that contain known phosphorylation and acetylation sites, as well as RNA-binding proteins. These proteins also displayed concerted subcellular localisations with enrichments for nucleolus across all datasets and mitochodrion in particular contexts. This reinforces our interpretation that for proteins with non-sigmoid behaviour, we are observing a mixture of behaviours from different functional copies of those proteins. This motivates expansion of the TPP method to deconvolute these behaviours, for example phosphoTPP (Huang *et al*., 2019; Potel *et al*., 2020; Smith *et al*., 2020) and other PTMs. The RNA-binding behaviour could be examined with high-throughput RNA-protein enrichment methods (Queiroz *et al*., 2019) and further deconvolution could be obtained by combining TPP with spatial proteomics methods (Mulvey *et al*., 2017; Geladaki *et al*., 2019). Though we observed non-sigmoidal behaviour in all datasets, more proteins were found to deviate in data generated from live cells (as compared to cell extracts).

As mentioned before, protein thermal stability can be affected by compound binding, PTMs and protein complex formation. In addition, protein solubility in cells might be affected by PTMs and other treatment-dependent effects, and even by ATP levels. Similar to protein solubility, compound treatment and other perturbations may affect the extent to which a protein is extracted in the applied experimental conditions leading to temperature dependent and temperature independent components that manifest themselves in thermal denaturation profiles. Whilst most referenced studies have been directed at identifying direct targets of small molecule inhibitors in live cells or in cell extracts, there is an increasing recognition of the potential of TPP as a methodology to profile molecular phenotypes (e.g. (Justice *et al*., 2020)) as it integrates multiple dimensions of regulation on proteome level into a single analytical approach. Such phenotyping could not only be informative for compound mechanism of action studies and to detect opportunities for combination treatments, but also to study effects of gene deletions, genetic variants and external stimuli and combinations there-off. As a consequence proteins can be affected in multiple ways and in different sub-cellular compartments resulting in more complex thermal denaturation behaviour than what can be robustly assessed with established computational approaches.

As demonstrated above our semi-parametric Bayesian approach is sensitive to detect protein effects that do not strictly follow the thermal denaturation-induced aggregation expected from isolated proteins and uniquely adds by identifying proteins affected by multiple parameters at once. Whilst not without challenges, the careful analysis of features in complex thermal denaturation curves is expected not only to facilitate hit calling but also to inform causality. This will be subject of future directions of our approach.

There are potential extensions of our methods to other TPP-based experimental designs (Mateus *et al*., 2020), to simultaneous joint modelling of multiple organisms (Jarzab *et al*., 2020) and to include prior information derived from other experiments. We could also use expected gain in information to optimise the drug concentration and temperatures used in the TPP experiments (Chaloner and Verdinelli, 1995). Summarising and normalisation to protein-level could also be avoided by modelling the data at (peptide spectrum match) PSM level. We have also used a default global prior for the prior model probabilities - these might be better specified using known prior properties about the drug being used.

As with all methods, our approach is not without limitations, for example increased computational cost could be a burden. However, if we are willing to sacrifice uncertainty quantification, we could simply use optimisation based inference instead. Our implementation is extensible with prior and model components easily change within our stan implementation. A Bioconductor package is in preparation (Huber *et al*., 2015).

## 4 Methods

### 4.1 Non-parametric analysis of response curves

We briefly describe the NPARC method for completeness (Childs *et al*., 2019). Let *y_ijk_* be the relative solubility of protein *i* at temperature *T_j_* for replicate measurement *k*. The null hypothesis states that the relative solubility of protein *i* at temperature *T_j_* is modelled as a single mean function regardless of the treatment condition or context:

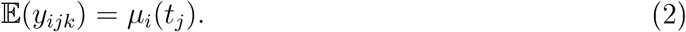

The alternative model allows for treatment effects or the mean function to change for each context

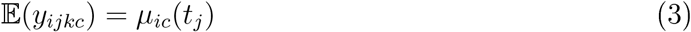

where *c* denotes the context. The mean function is modelled using the 3-parameter sigmoid model:

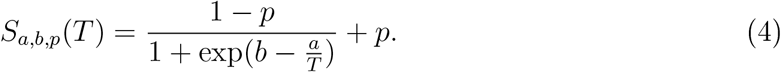

To clarify, under *H*_0_ the parameters *a,b,p* are fixed for both contexts, whilst under the alternative *H*_1_ the parameters *a,b,p* are allowed to be context specific. For hypothesis testing, the F statistic is computed

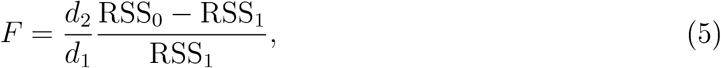

where *RSS*_0/1_ denotes the sum of the squared residuals when fitting the null (0) or the alternative (1) model and *d*_1/2_ are referred to as degrees of freedom. Large values of the *F* statistic represents reproducible changes thermal stability. If the residuals were *i.i.d* normally distribution then we could perform an *F*-test using the null distribution *F*(*d*_1_, *d*_2_), where the degrees of freedom are computed from simple parameter and observation counting. However, the *i.i.d* assumption do not hold and so Childs *et al*. (2019) estimate the null distribution using new *effective* degrees of freedom 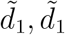. Approximating the null distribution assumes a unimodal null distribution and that the majority of observations are samples from the null distribution. We refer to Childs *et al*. (2019) for detailed formulae. Once the approximate null has been obtained *p*-values can be computed as usual and multiple hypothesis testing correction applied (Benjamini and Hochberg, 1995).

### 4.2 Bayesian inference and model selection

#### 4.2.1 Bayes’ theorem and hypothesis testing

In this section, we summarise Bayesian inference and model selection. The advantage of the Bayesian framework is that we no longer need to estimate a null distribution and multiplicity is automatically controlled via the prior model probabilities. This avoids making any assumptions about the properties of the null distribution. Furthermore, prior information is included on the parameters, which has a number of benefits, including allowing the shrinkage of residuals towards 0, regularising the inferred parameters and improving algorithmic stability. Furthermore, in a Bayesian analysis, we obtain samples from the posterior distribution of the parameters and hence the posterior distribution of the mean function can be obtained to quantify uncertainty.

Bayesian inference begins with a statistical model 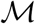 of the observed data *y* with the parameters of the model denoted by *θ*. Given a prior distribution for the parameters, denoted 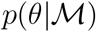, and observed data *y*, Bayes’ theorem tells us we can update the prior distribution to obtain the posterior distribution using the following formula:

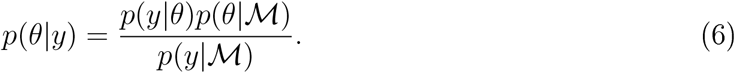

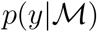 is referred to as the marginal likelihood, since it is obtained by marginalising *θ*:

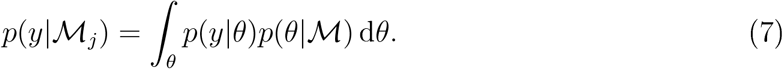

The task of hypothesis testing can be cast as a model selection problem. Indeed, the null hypothesis is associated with a model 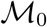, whilst the alternative hypothesis is associated with a model 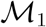. Thus, the task of hypothesis testing is that of selecting between two competing models.

To perform model selection, we are interested in the following posterior quantity (Berger and Molina, 2005),

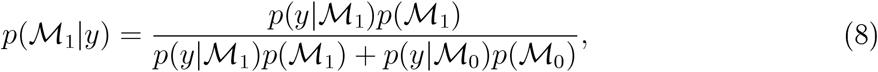

that is the posterior model probability, given the data. The relative plausibility of two model is quantified through the posterior odds, which is the prior odds multiplied by the Bayes factor (Kass and Raftery, 1995).

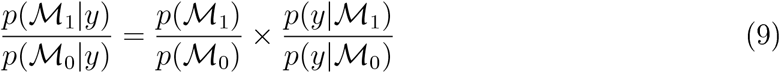

The challenging of computing these equations is obtaining the marginal likelihood (equation 7). We note that because of the integration with respect to the prior there is automatic penalisation of additional model complexity. The marginal likelihood is challenging to compute and is only available in analytic form for a small number of relatively simple models.

A number of sampling based approach are available to compute marginal likelihoods, such as bridge sampling (Meng and Wong, 1996; Meng and Schilling, 2002), path sampling (Gelman and Meng, 1998), importance sampling (Robert and Wraith, 2009), harmonic mean sampling (Gelfand and Dey, 1994), nested sampling (Skilling *et al*., 2006; Chopin and Robert, 2010; Johnson *et al*., 2015) (see also (Carlin and Chib, 1995)). Though these sampling based approaches produce highly accurate marginal likelihoods, these approaches require excessive computation in our case. Instead, we approximate the marginal likelihood using the Metropolis-Laplace estimator. Briefly, the log of the marginal likelihood (equation 7) is estimated as (Lewis and Raftery, 1997):

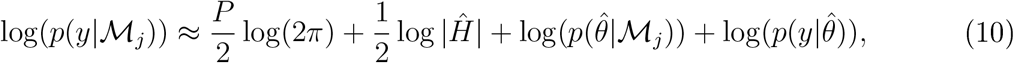

where 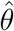 a Monte-Carlo estimator of the parameters, *P* is the number of parameters and 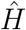 is estimated by the sample covariance of the posterior samples. This approach is used for both the Bayesian sigmoid model and the semi-parametric model.

Finally, we have yet to specify the prior model probabilities *p*(*M_j_*) for *j* = 0,1. To control for multiplicity, we can adjust the prior model properties to assume that the null model is more likely that the alternative (Scott and Berger, 2006). Hence, we set 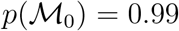 and 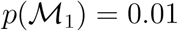.

#### 4.2.2 Posterior predictive checks and out-of-sample predictive performance

Formal model selection via the marginal likelihood can be used to select between two or more competing models. However, models can also be assessed and criticised using measures of predictive performance. Here, we consider posterior predictive checks, as well as out-of-sample predictive performance. A posterior predictive check begins with simulating from the posterior predictive distribution:

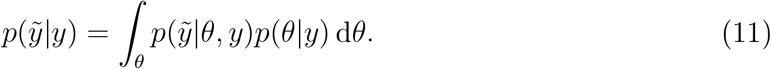

This is the distribution obtain by marginalising the distribution of 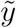 given *θ* over the *posterior* distribution of *θ* given *y*. The rationale is that simulated data from the posterior predictive should look similar to the observed data (Gelman *et al*., 2013). We simulate these datasets *y*_rep_ and compute the 50% and 95% credible bands, for the models of interest. Though other posterior predictive summaries can be used, such as Kernel Density Estimate posterior predictive checks (see supplement).

Another approach is to examine the out-of-sample predictive accuracy from the fitted Bayesian models. We use (approximate) leave-one-out cross validation (LOO-CV) with the log predictive density as the utility function (equivalently the log-loss) (Vehtari *et al*., 2017):

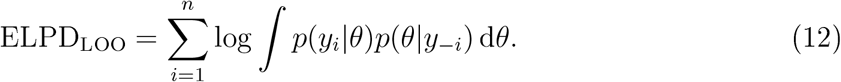

Equation 12 is the leave-one-out predictive density given the observed data without the *i^th^* observation, summed over the observations. This process is intensive so the expected log pointwise predictive density (ELPD) is estimated using Pareto smoothed importance sampling (PSIS) (Vehtari *et al*., 2017).

### 4.3 Bayesian sigmoid model

In this section, we develop our Bayesian sigmoid model. For our proposed Bayesian sigmoid model, we assume the aforementioned sigmoid model. As before, under 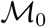 a single sigmoid model is posited irrespective of any treatment effects or contexts. While the competing model 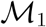 allows the sigmoid parameters to be context specific. Thus under the null hypothesis, we assume

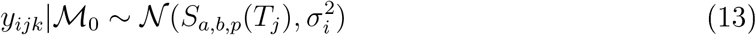

whilst for the competing model

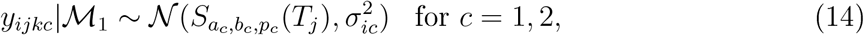

where again c denotes the context or treatment effect. To complete the specification of our model, we need to declare the priors. The sigmoid shape parameters *a, b* are required to be positive and thus we place a Gamma distribution on these parameters. The right tail of the Gamma distribution discourages posterior mass on excessively large values of *a* and *b*. To obtain reasonable defaults for these priors, we examined the fitted values found by previous analysis (Childs *et al*., 2019), as well as performing a prior predictive check (Box, 1980). Thus priors are specified for *a,b* as follows

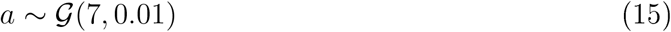

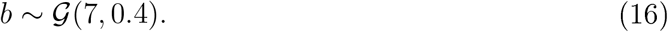

The parameter *p* is restricted between 0 and 1 and thus a Beta prior is appropriate for this parameter. Given that the plateau is generally close to 0 and rarely above 0.5 we specify the following prior

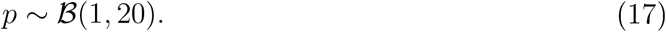

For the standard deviation of the residuals *σ*, we desire these to be considerably smaller than the scale of the data and shrunk towards 0. This has two purposes: the first is that we want the data to be explained by variations in the mean function not simply by wide errors; secondly smaller residuals allow us to discriminate between small but reproducible shifts in melting profiles. We opt for the folded-normal distribution on σ (Gelman *et al*., 2006). We specify the prior as follows

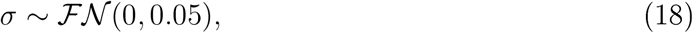

which puts significant mass around 0 to encourage shrinkage, whilst residuals up to 0.4 are not considered surprising. There is no conjugacy between our prior and likelihood, which makes obtaining samples from the posterior distribution challenging. We employ Hamiltonian Monte-Carlo (Duane *et al*., 1987), in particular a variant of the no-u-turn sampler (Hoffman and Gelman, 2014; Betancourt, 2017) with an implementation in Stan (Bürkner *et al*., 2017; Carpenter *et al*., 2017).

### 4.4 Bayesian semi-parametric model

Our Bayesian sigmoid model allowed us to remove the assumptions relating to the estimating the null distribution, but still assumes a sigmoid functional form and uncorrelated residuals. To relax these assumptions, we propose a semi-parametric model. We assume the parametric sigmoid function and introduce an additional term so that the melting curves for protein *i* are modelled according the following (suppressing notation on the condition)

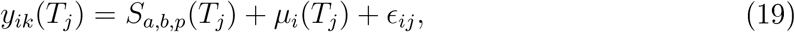

where *μ* is some deterministic function of temperature and 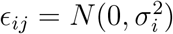 is a noise variable. Without any suitable parametric assumptions for *μ_i_*, we perform inferenee for *μ_i_* by specifying a *Gaussian process prior*, so that:

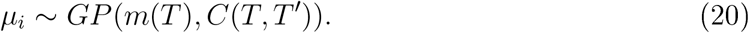

A Gaussian process (GP) prior is uniquely determined by its mean and covariance function, which determine the mean vectors and covariance matrices of the associated multivariate Gaussians. We do not have any prior believe about the symmetry or periodicity of our functions (beyond what is already encoded by *S_a,b,p_*) and thus we specify a centred GP with a squared exponential covariance function

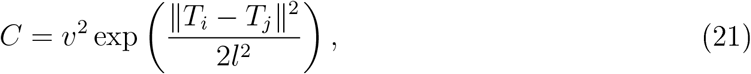

where *v*^2^ is a marginal variance parameter and *l*, a length-scale parameter, encodes the distance at which observations are correlated. The adopted GP prior of *μ_i_* tells us that the relative solubility for protein *i* is modelled as follows

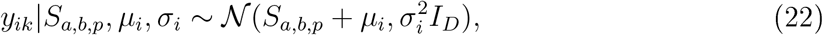

where *D* denotes the number of measured temperatures. Note that we can make *n_i_* repeated measurement (or replicates) of protein *i* at temperature *T_j_*. We denote *y_i_* = {*y_i1_*, ..,*y_in_i__*} to be the concatenation of replicate measurements. Hence, the above implies that

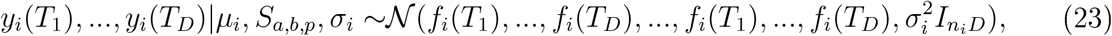

where *f_i_*(*T*_1_),..., *f_i_*(*T_D_*) is repeated *n_i_* times and *f_i_*(*T_j_*) = *S_a,b,p_*(*T_j_*) + *μ_i_*(*T_j_*). Our GP prior tell us that

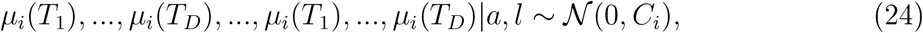

where *C_i_* is an *n_i_D* × *n_i_D* matrix. Note that the above means that we can marginalise *μ_i_* to avoid inference of this unknown function and obtain:

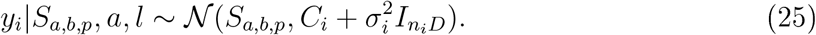

Reintroducing the context or treatment effect, we allow the parameters to vary between them. Thus, under the null hypothesis, we assume

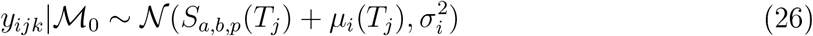

whilst for the competing model

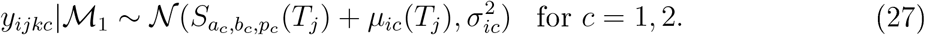

To complete our model, we need to specify the prior distributions. For parameters in common with the sigmoid model we make the same prior choices. Thus, it remains to make prior choices for *v* and *l*. The challenges of specifying priors for the hyperparameters of the Gaussian process are well documented (Berger *et al*., 2001; Paulo *et al*., 2005; De Oliveira, 2007; van der Vaart *et al*., 2009; Fuglstad *et al*., 2019). To obtain a sensible prior it is important to note that our model is weakly non-identifiable. This is because the non-parametric part can explain the parametric components. However, this is not, in general, an issue for Bayesian analysis. To advert problems this can cause for inference, we have to make judicious prior choices.

The first step is to encourage the marginal variance parameter to be on the scale of the residuals rather than that of the data. We already placed a folded-normal prior on the measurement error *σ*. For the marginal variance *v*^2^, we impose even stronger shrinkage towards 0 by using a folded-student-t prior. This prior also has heavy tails allowing the non-parametric term to explain complex variations, if supported by the data. To summarise, we specify

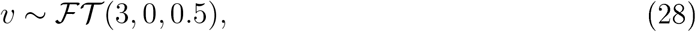

where 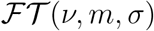 denotes a folded-student-t density with degrees of freedom *ν*, mean *m* and scale *σ*. On the other hand, for the length scale parameter *l*, we wish to avoid excessively small values. Short length-scales allow the Gaussian process simply to interpolate the data and overfit. Thus, we propose a log-normal prior for l, which has a sharp left tail and heavy right tail, discouraging small length scales and really large length scales, respectively. We find that the following prior works well in practice (sensitivity is tested in the supplement):

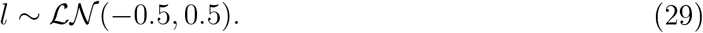

Inference for Bayesian models that incorporate Gaussian processes priors can be computationally intensive and so we make use of reduced-rank Gaussian process methods by approximating the covariance function (Solin and Särkkä, 2020). As with the sigmoid model our semi-parametric model is implemented in Stan (Carpenter *et al*., 2017).

## 6 Acknowledgements

We thank members of the Cambridge Centre for Proteomics and Nils Kurzawa for insightful discussions.

## Appendix 1 : Residual correlation matrices for TPP datasets

### Panobinostat residual correlation matrix

**Figure 7:**
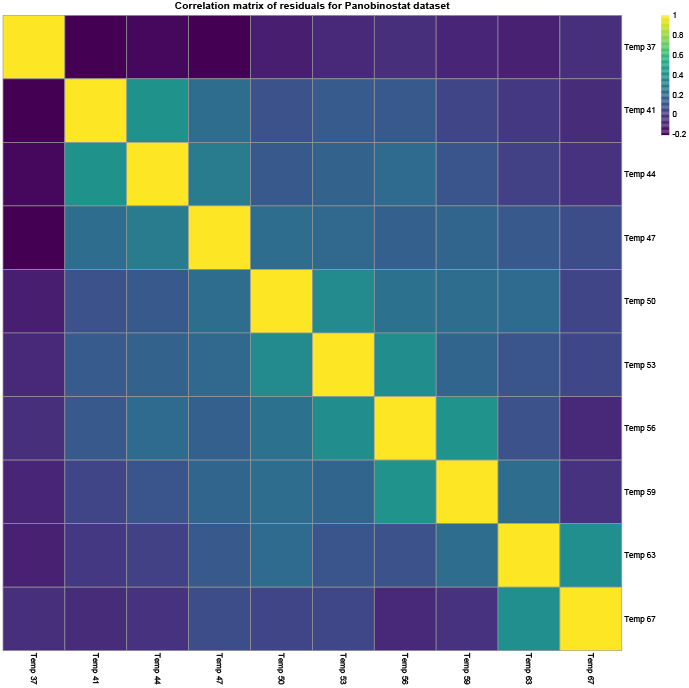
Sample Spearman correlation matrix for the residuals from the sigmoid model applied to Panobinostat dataset

### Dasatinib 0.5 residual correlation matrix

**Figure 8:**
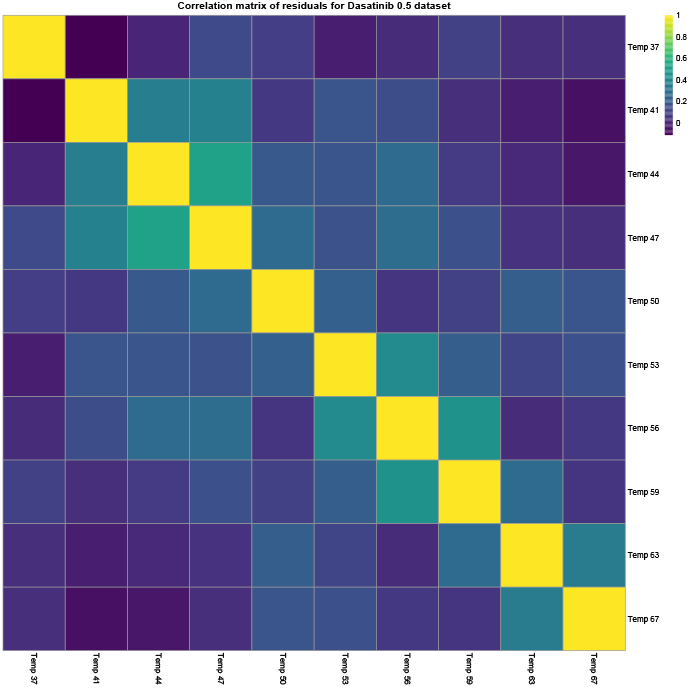
Sample Spearman correlation matrix for the residuals from the sigmoid model applied to Dasatinib 0.5 dataset

### Dasatinib 5 residual correlation matrix

**Figure 9:**
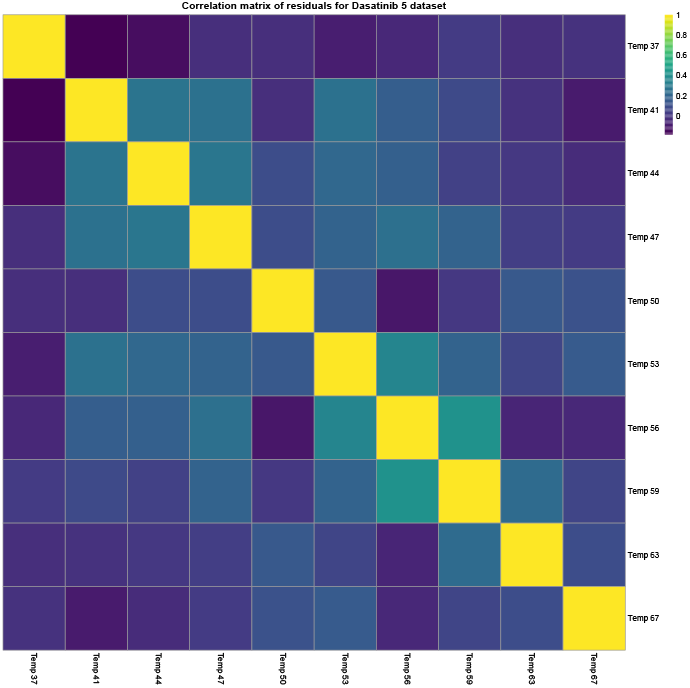
Sample Spearman correlation matrix for the residuals from the sigmoid model applied to Dasatinib 5 dataset

## Appendix 2 : Analysis of the ATP dataset

The ATP dataset as described in the main text was also analysed to determine the sensitivity of our approaches. In this case, uniprot key-terms for “ATP-binding” were extracted and used as a pseudo ground truth. We then compute the sensitivity for each of the approaches at threshold *p* = 0.01 or posterior probability at 0.99. It is clear that the Bayesian approaches outperform the NPARC approach in this case.

**Figure 10:**
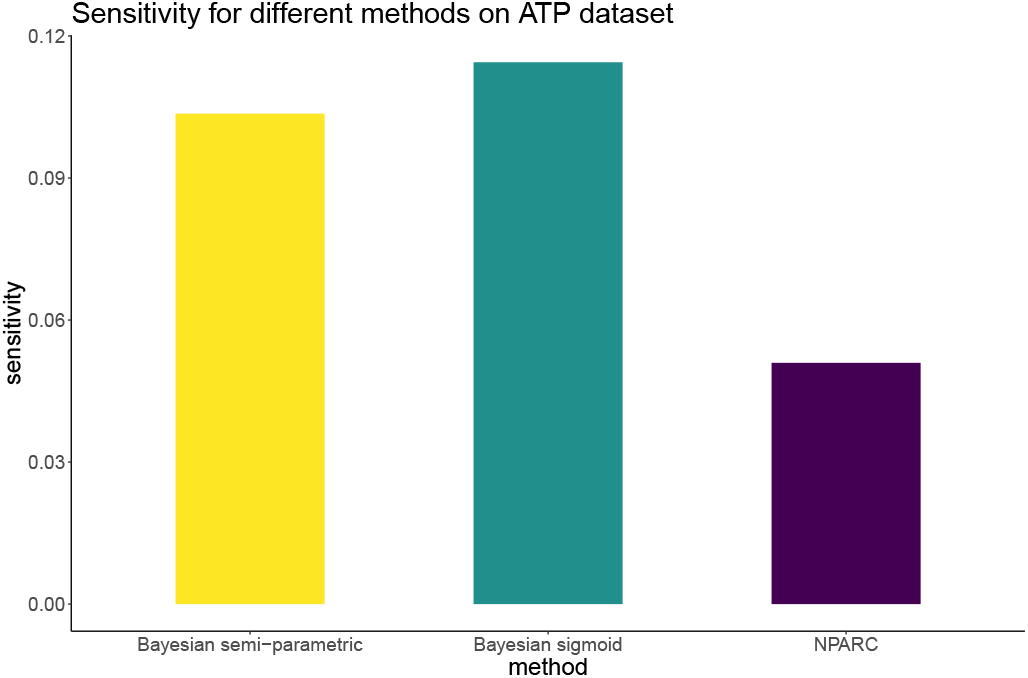
Sensitivities for the different methods

## Appendix 3 : Simulation specification

To characterise the methods when there is a known ground truth, we perform a simulation study.

### Simulation 1

Here we describe the first simulation study. We let *T* = (35, 40, 45, 50, 55, 60, 65). The first 249 proteins are simulated form the following model, where there is no difference between the control and treatment context. Hence, for each protein *i*, 4 protein profiles are drawn for the following distribution

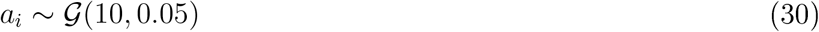

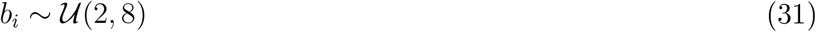

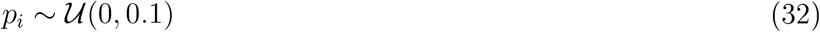

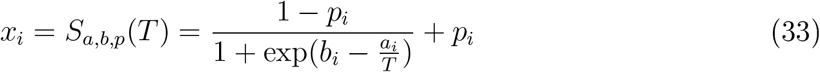

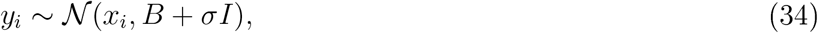

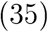

where *σ* = 0.01 and *B*(*i,j*) = exp(−0.4(*T_i_* − *T_j_*)^2^) for *i* ≠ *j* and 0 otherwise. The next 201 proteins are drawn from the following distribution, again with no difference between control and treatment, with 4 profiles drawn for each protein.

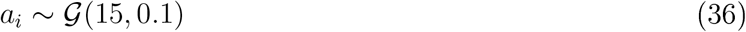

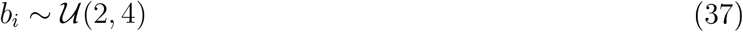

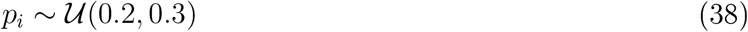

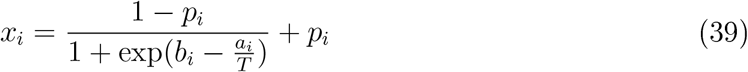

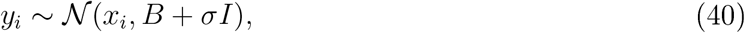

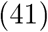

where *σ* and *B* are as above. The above scenario is to designed to generate a null sampling population that is a mixture distribution. For cases where the treatment does have an effect on the melting profile, we simulate in the following way. For each protein *i* and each context *c* = 1, 2 simulate the following:

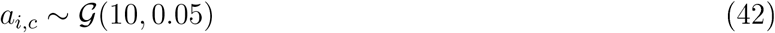

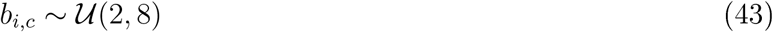

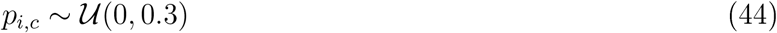

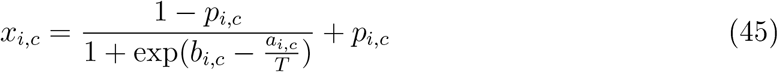

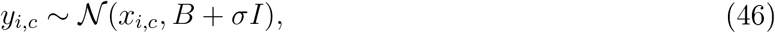

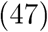

where *σ* = 0.001, for a total of 25 protein with 2 replicates for each context. The remaining 25 proteins are drawn from the following

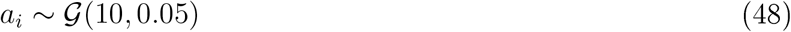

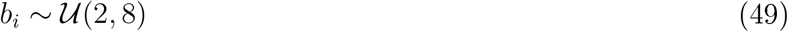

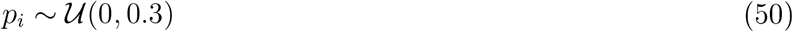

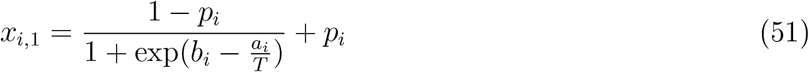

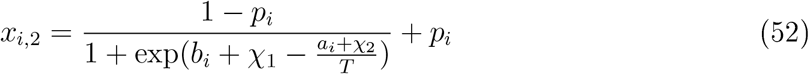

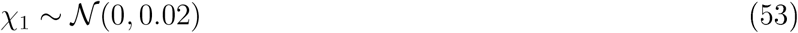

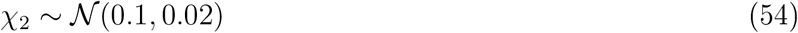

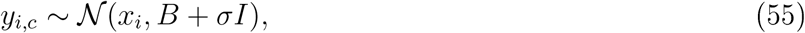

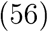

where *σ* = 0.001. These proteins are drawn using a different mechanism to allow flexibility in the ways the context effects the protein profiles. This simulation is designed such that the majority of proteins are drawn from the null sampling distribution; however, there are two mechanism to generate the null and alternative models. This violate the assumptions of the NPARC model, we also note that the models are mis-specified with respect to the Bayesian models as well.

### Simulation 2

Here we describe the second simulation study and we let *T* = (35, 40, 45, 50, 55, 60, 65). The first 249 proteins are simulated form the following model, where there is no difference between the control and treatment context. Hence, for each protein *i*, 4 protein profiles are drawn for the following distribution

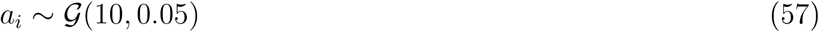

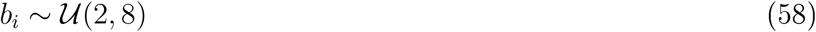

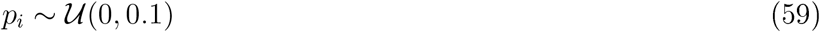

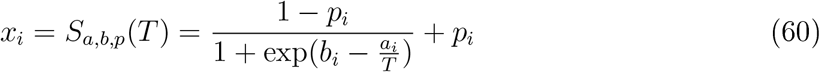

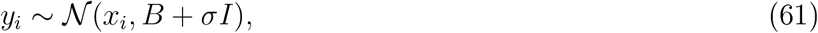

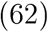

where *σ* = 0.01 and *B*(*i,j*) = exp(−0.4(*T_i_* − *T_j_*)^2^) for *i* ≠ *j* and 0 otherwise. For the next 101 proteins are drawn from the following, again with no difference between control and treatment, with 4 profiles drawn for each protein.

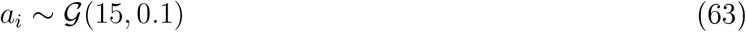

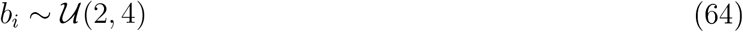

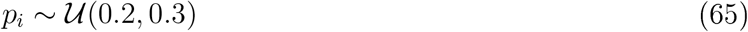

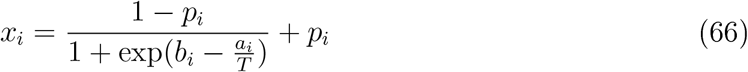

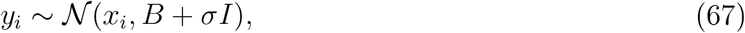

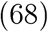

where *σ* and *B* are as above. The above scenario is to designed to generate a null sampling population that is a mixture distribution. For cases where the treatment does have an effect on the melting profile, we simulate in the following way. For each protein *i* and each context *c* = 1, 2 simulate the following:

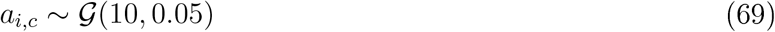

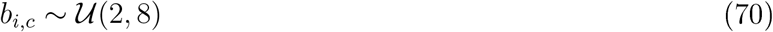

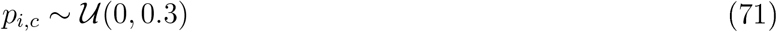

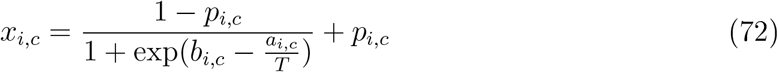

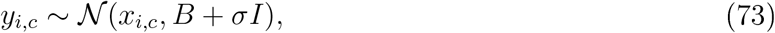

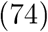

where *σ* = 0.001, for a total of 100 protein with 2 replicates for each context. The remaining 50 proteins are drawn from the following

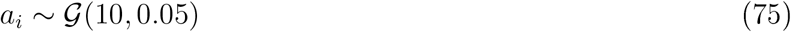

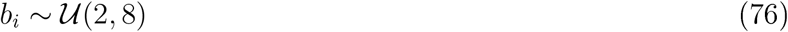

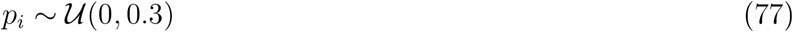

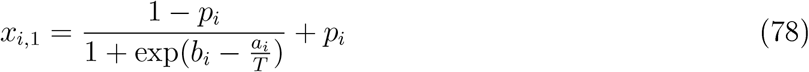

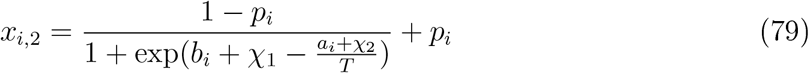

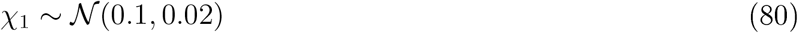

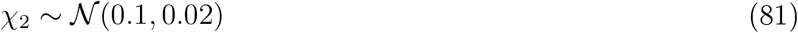

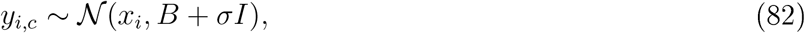

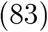

where *σ* = 0.001. These proteins are drawn using a different mechanism to allow flexibility in the ways the context effects the protein profiles. This simulation is designed such that only 70% of proteins are drawn from the null sampling distribution and there are two mechanism to generate the null and alternative models. These violate the assumptions of the NPARC model, we also note that the models are mis-specified with respect to the Bayesian models as well.

### Simulation 3

The same for simulation study 1, but the prior on the length scale is change to 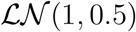 to test the sensitivity of the analysis.

## Appendix 4: Simulation study results

This section reports the simulation study results. All approaches are capable of controlling the FDR and, as we observed for real data, the Bayesian models show improved sensitivity.

Since the simulation temperatures are at different increments to the real data, the default priors are changed to 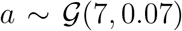 and 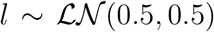. The other priors are held the same.

### Simulation 1

The simulation study 1 is performed 10 times to report a distribution of results. We report the distribution of sensitivity at a fixed specificity of 0.99 for all methods (figure 11). We also plot an example ROC curve from these simulations in figure 12. It is clear that the Bayesian models shows improvement over NPARC and that the semi-parametric model is an improvement over the sigmoid models.

**Figure 11:**
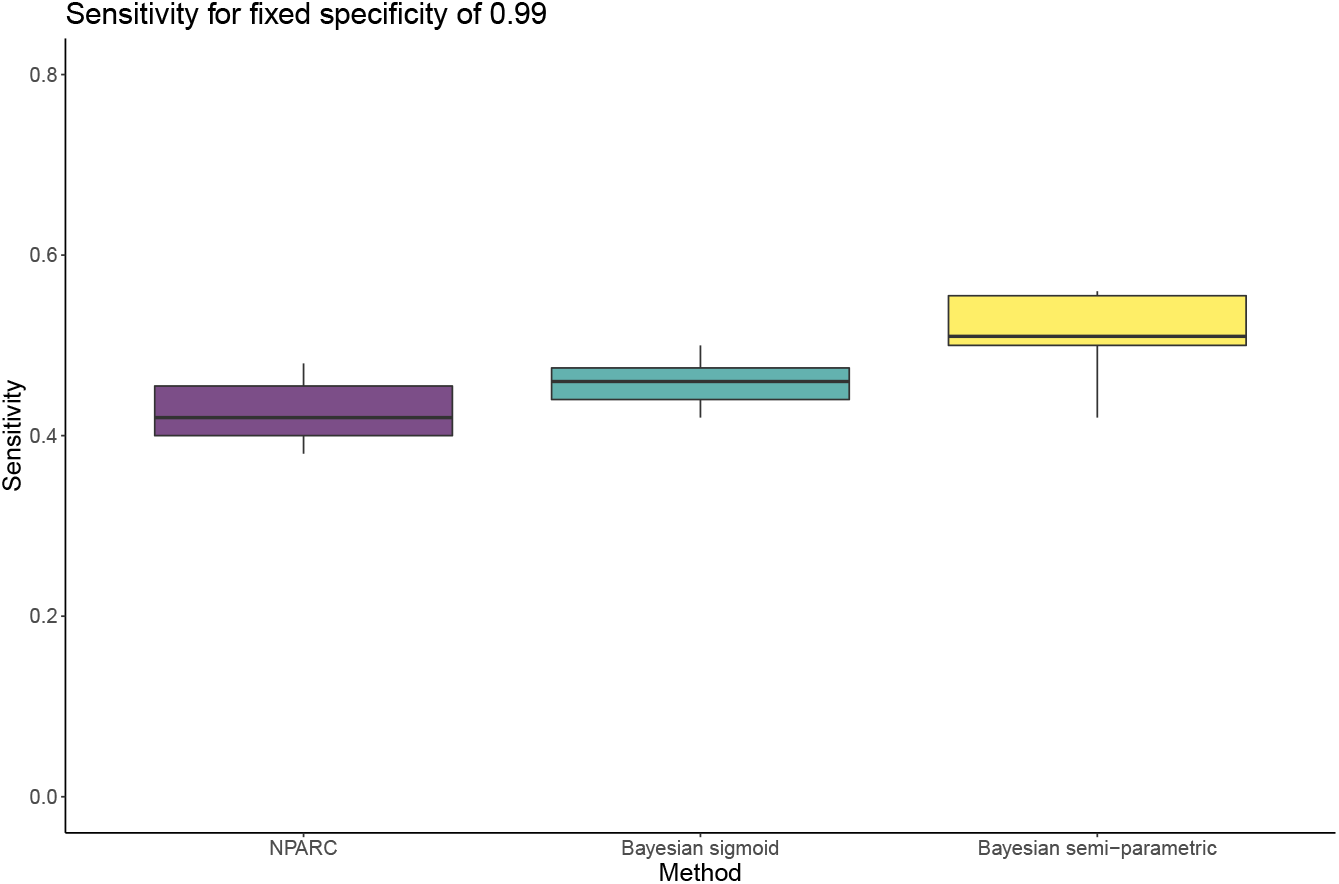
Sensitivities for a fixed specificity of 0.99. The boxplots are distributions from repeated simulations.

**Figure 12:**
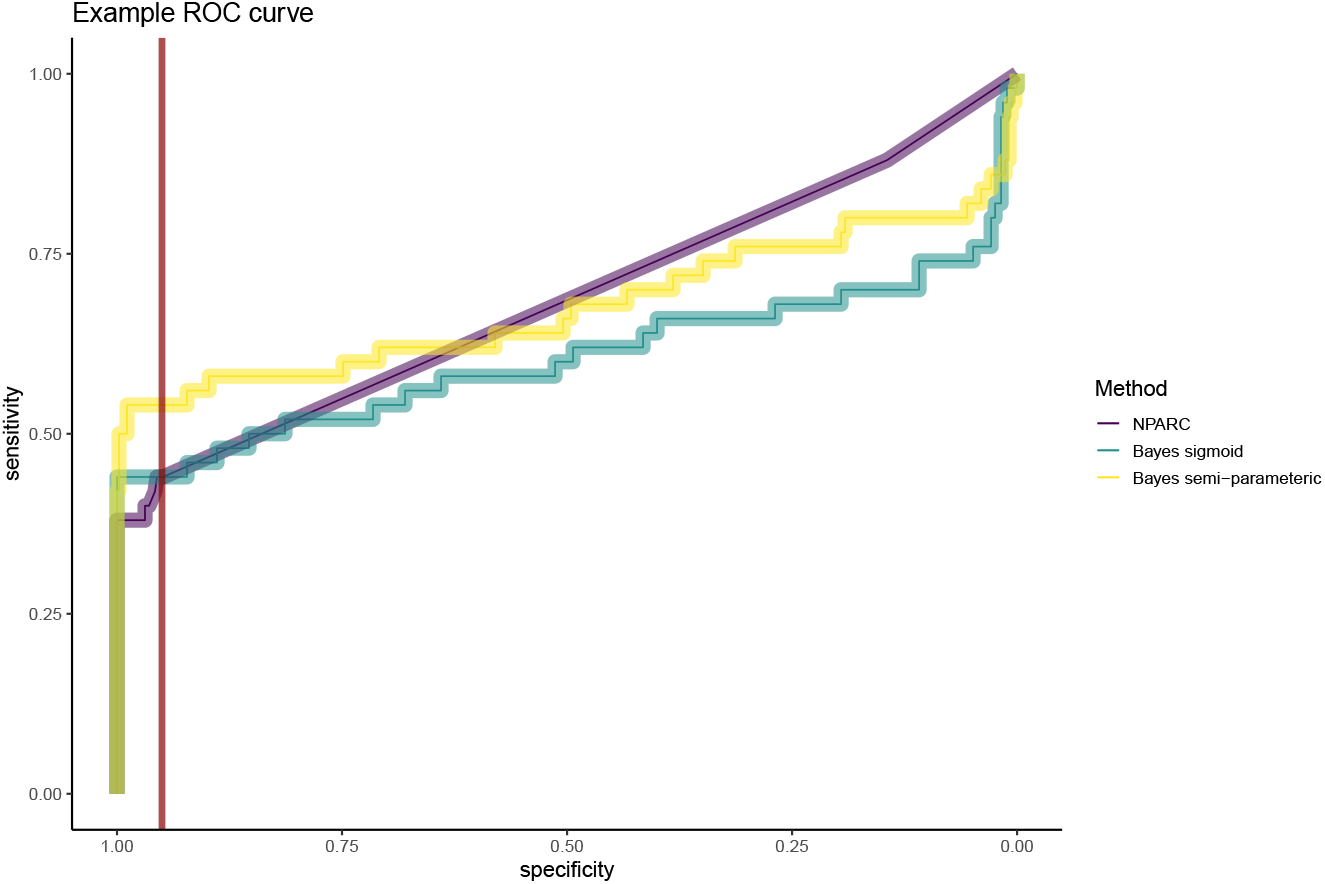
An example ROC curve from the simulations. The dark red line indicates a specificity of 0.95

### Simulation 2

The simulation study 2 is performed 10 times to report a distribution of results. We report the distribution of sensitivity at a fixed specificity of 0.99 for all methods (figure 13). We also plot an example ROC curve from these simulations in figure 14. It is clear that the Bayesian models shows improvement over NPARC and that the semi-parametric model is an improvement over the sigmoid models.

**Figure 13:**
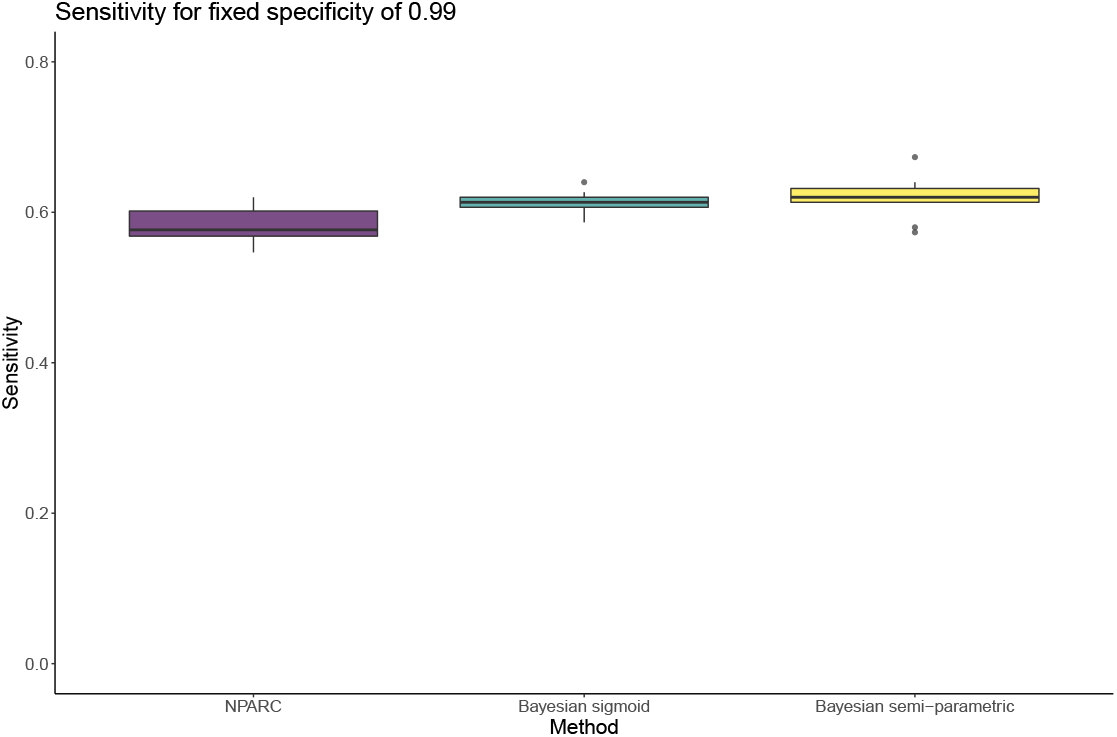
Sensitivities for a fixed specificity of 0.99. The boxplots are distributions from repeated simulations.

**Figure 14:**
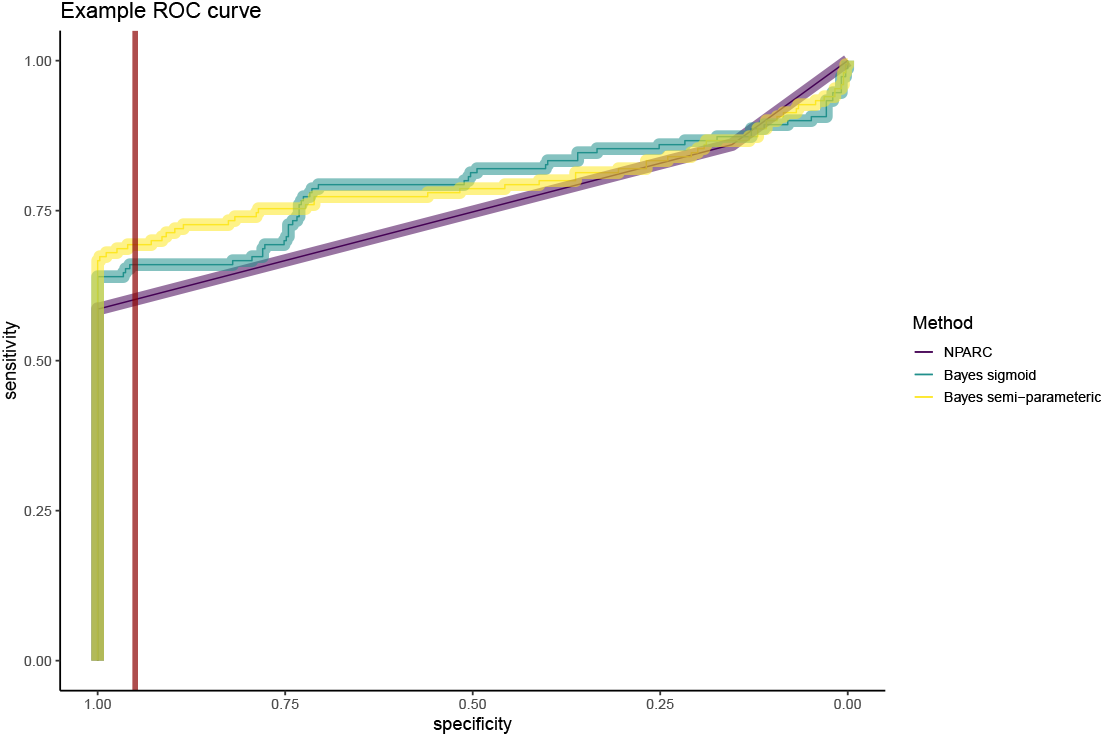
An example ROC curve from the simulations. The dark red line indicates a specificity of 0.95

### Simulation 3

The simulation study 3 is performed 10 times to report a distribution of results. We report the distribution of sensitivity at a fixed specificity of 0.99 for all methods (figure 15). We also plot an example ROC curve from these simulations in figure 16. It is clear that the Bayesian models shows improvement over NPARC and that the semi-parametric model is an improvement over the sigmoid models.

**Figure 15:**
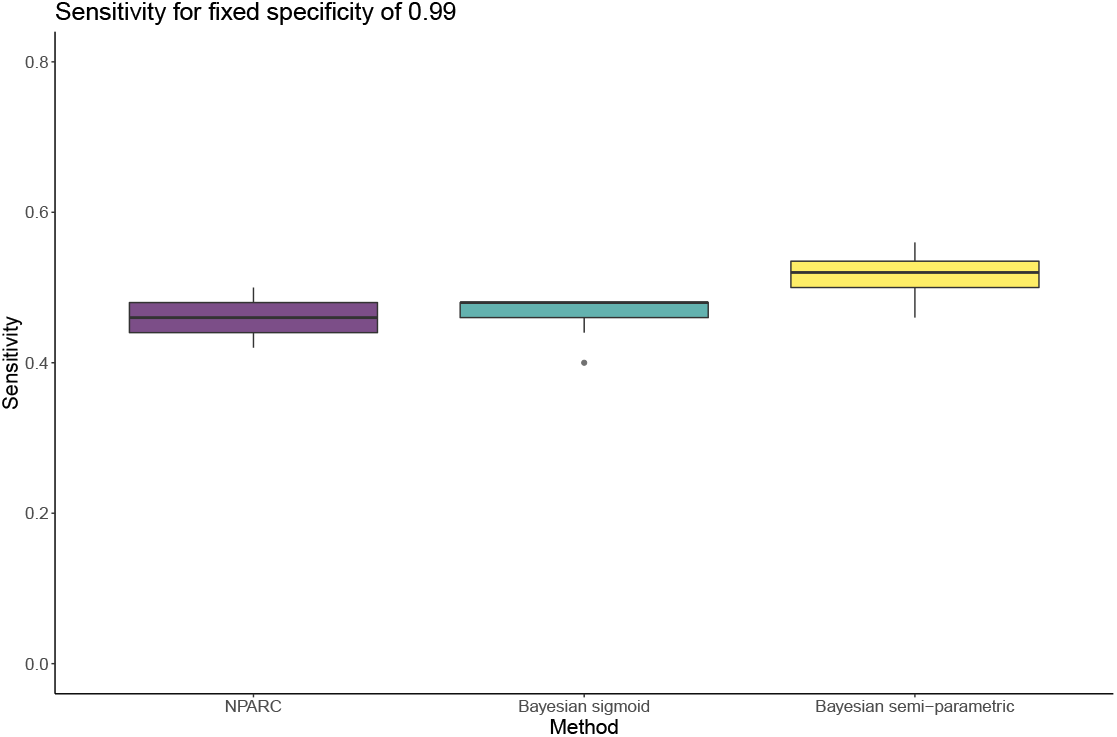
Sensitivities for a fixed specificity of 0.99. The boxplots are distributions from repeated simulations.

**Figure 16:**
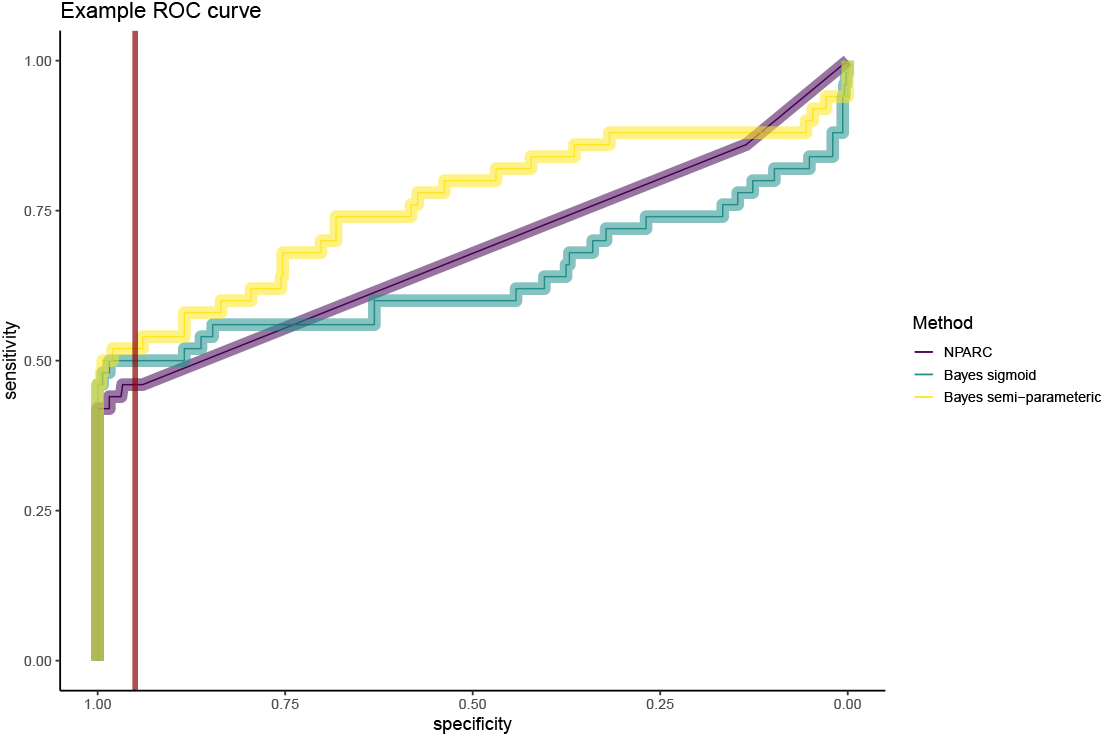
An example ROC curve from the simulations. The dark red line indicates a specificity of 0.95

## Appendix 5: Model selection

In this section, we further evaluate the Bayesian sigmoid model and Bayesian semi-parametric approach. We examine the marginal likelihood, averaged over proteins and report the distribution over simulations. Figure 17 shows that the marginal likelihood for the semi-parametric is considerably higher than the Bayesian sigmoid model. We also evaluate out-of-sample predictive performance for both Bayesian models. We examine the leave-one-out cross-validation estimate with log scoring. Figure 18 shows that the Bayesian sigmoid model has higher expected log predictive density (averaged over proteins). This is perhaps antagonistic with the log marginal likelihood; however, it is well documented that non-parametric models can reduce out-of-sample predictive performance (Powell, 1994). That being said the improvement is less than 2 points.

**Figure 17:**
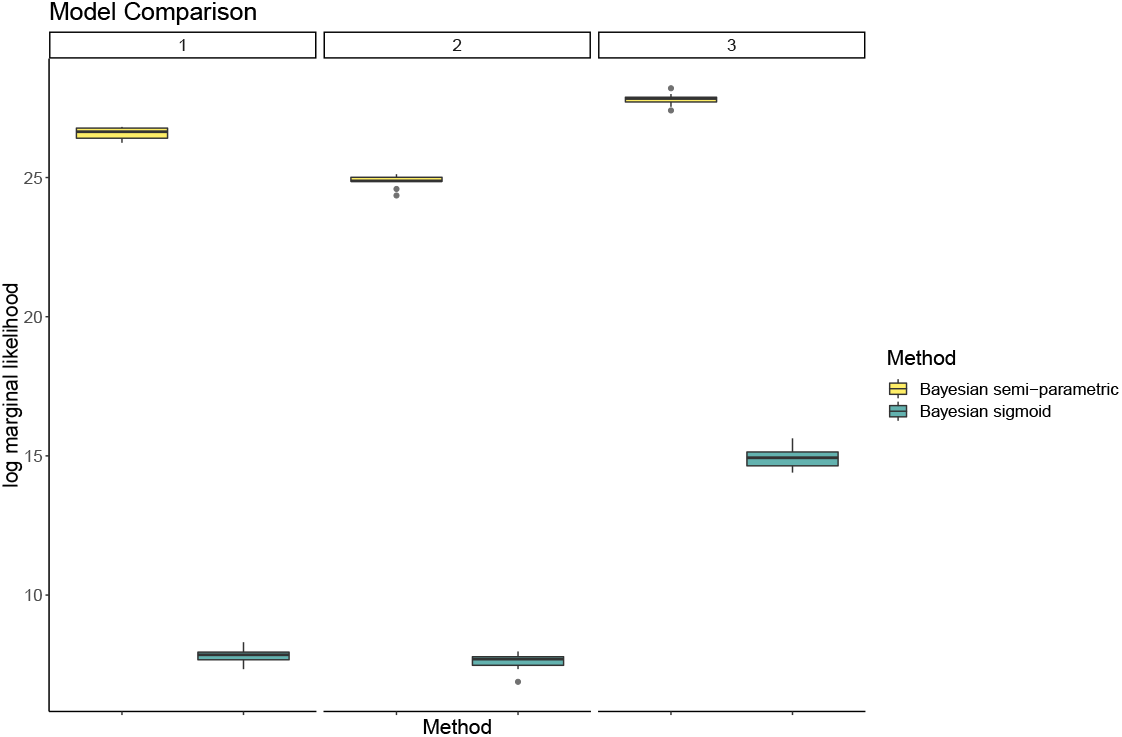
Marginal likelihoods for the different Bayesian models where distributions are over different simulations.

**Figure 18:**
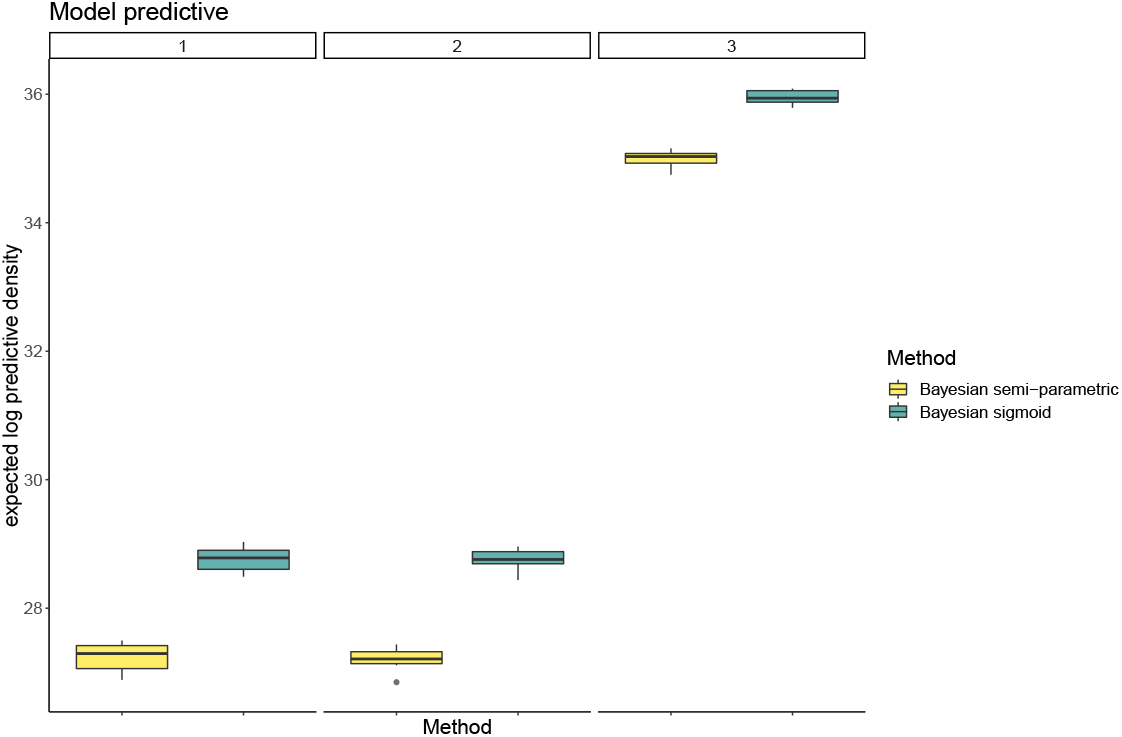
LOO-CV estimates of the expected log predictive density, averaged over proteins. The distributions are over different simulations

## Appendix 6: Robustness to model miss-specification

In this section, we test robustness of the method to model miss-specification. To be more precise, we consider a simulation scenario where the model is not from the class of models being fit. Recall simulation scenario 1 and instead of simulating data from the 3-parameter sigmoid model, we simulate for the following 4-parameter sigmoid model:

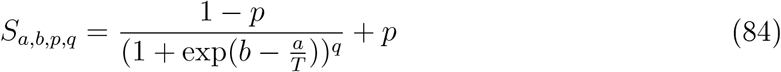

Note that in this 4-parameter sigmoid model, a degree of asymmetry is allowed. We set *q* = 10 for this simulation and apply NPARC and the Bayesian model. The simulations are performed 10 times to generate a distribution of results. Fortunately, both the Bayesian models are robust to this model miss-specification and are able to successfully control the false positive rate and have high sensitivity (see figures 19 and 20). NPARC is much more sensitive to model miss-specification.

**Figure 19:**
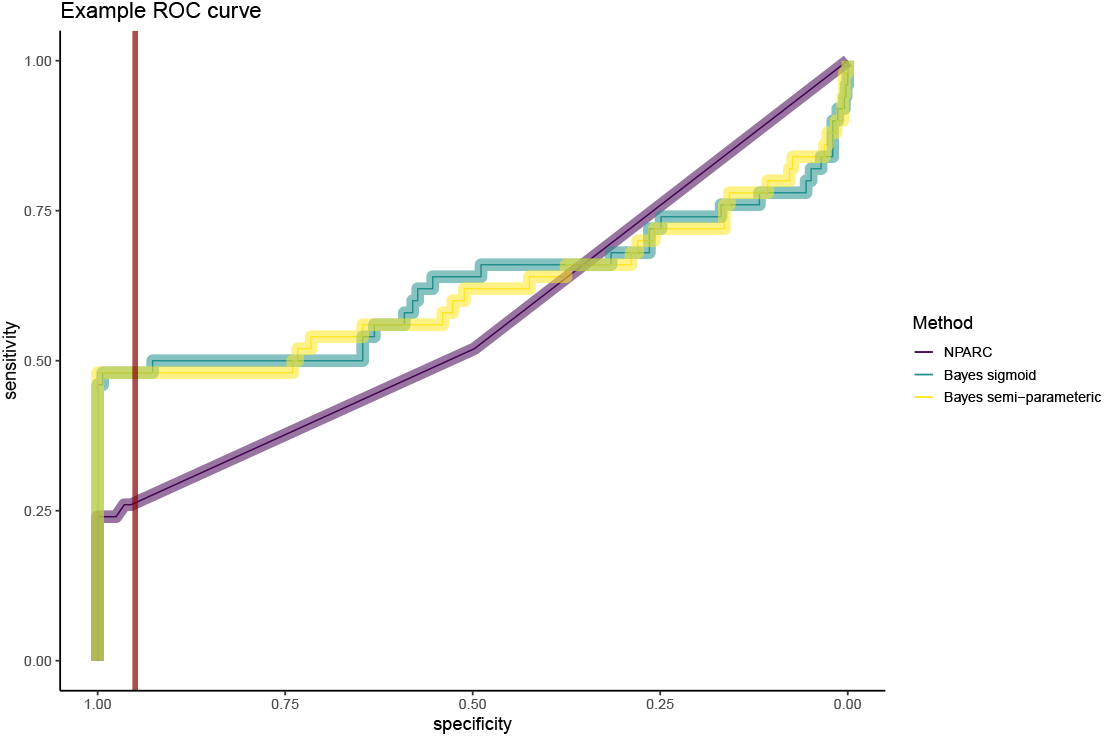
An example ROC curve from the miss specified model. The dark red line indicates a specificity of 0.95

**Figure 20:**
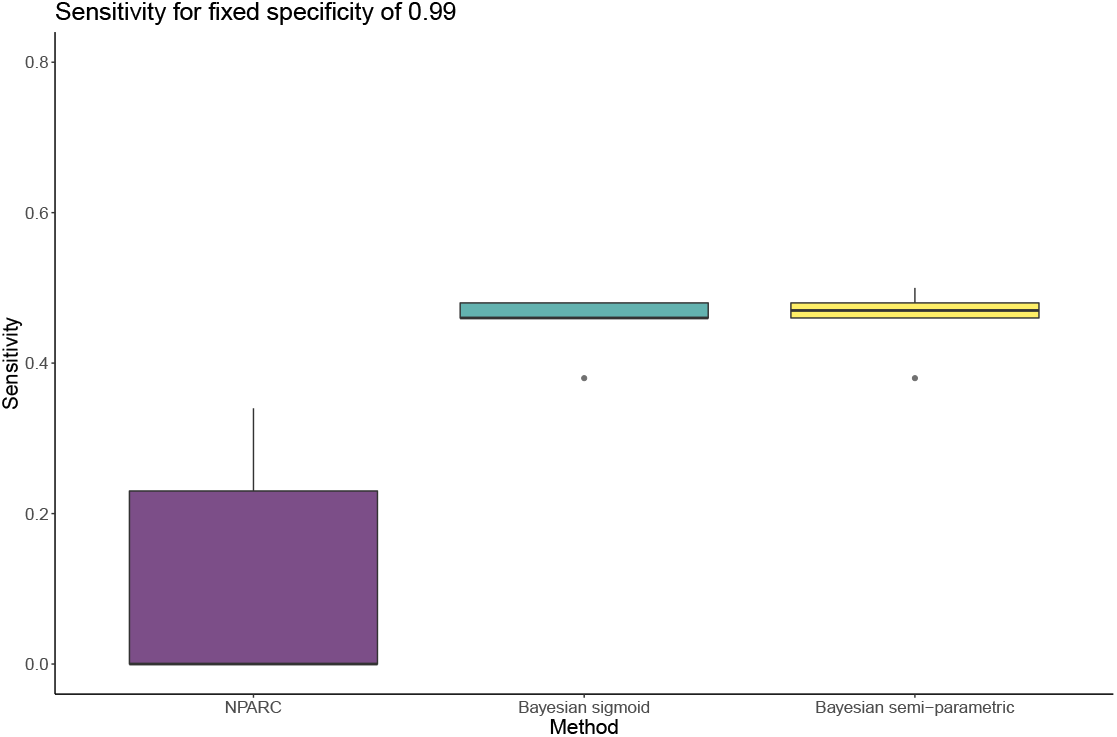
Sensitivities for a fixed specificity of 0.99 for miss specified model. The boxplots are distributions from repeated simulations.

## Appendix 7: Posterior predictive checks

To complement the posterior predictive checks in the main text for AP4S1. We simulate 100 datasets *y*_rep_ from the posterior predictive distribution of each of the two Bayesian models. For each dataset, we compute the kernel density estimate of the simulated datasets, as well as the observed data. We plot the results for the sigmoid model in figure 21 and the semi-parametric model in figure 22. It is evident that the posterior predictive check for the semi-parametric model is more appropriate for the observed data.

**Figure 21:**
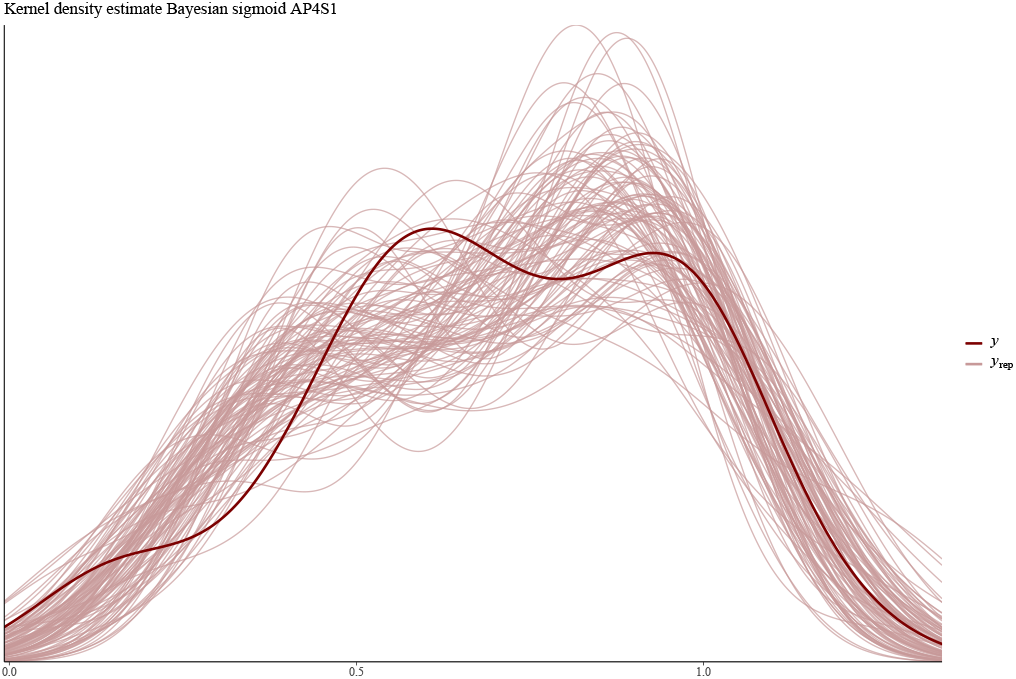
Kernel density estimate posterior predictive check for sigmoid model

**Figure 22:**
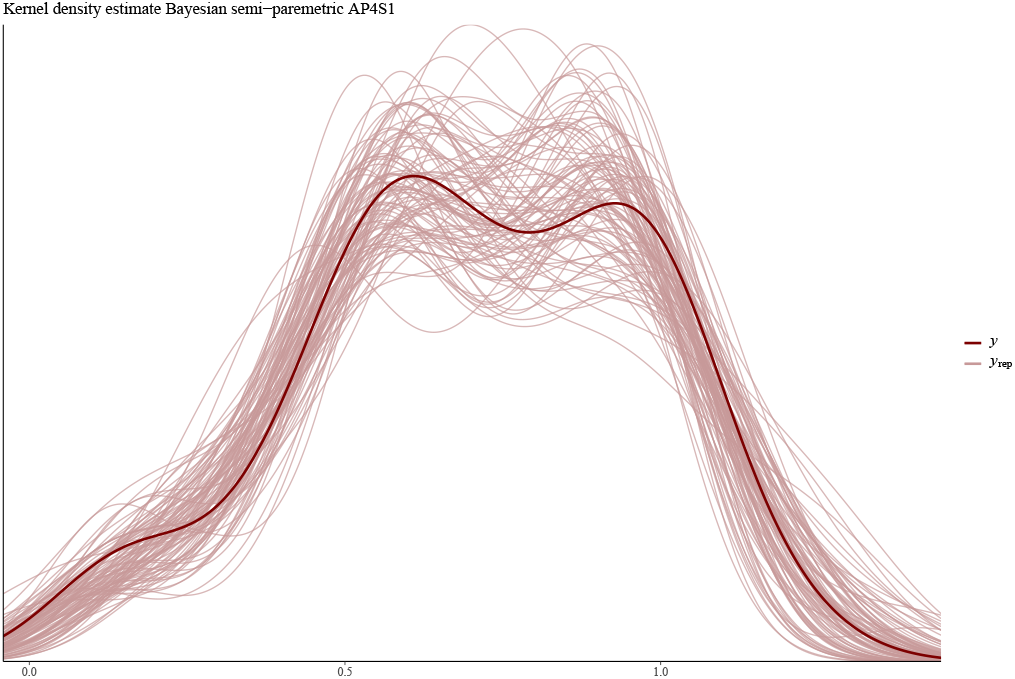
Kernel density estimate posterior predictive check for semi-parametric model

## References

Allis, C. D. et al. (2016). The molecular hallmarks of epigenetic control. Nature Reviews Genetics, 17(8),487.

Babtie, A. C. et al. (2014). Topological sensitivity analysis for systems biology. Proceedings of the National Academy of Sciences, 111(52), 18507–18512.

Ball, K. A. et al. (2020). An isothermal shift assay for proteome scale drug-target identification. Communications biology, 3(1), 1–10.

Becher, I. et al. (2016). Thermal profiling reveals phenylalanine hydroxylase as an off-target of panobinostat. Nature chemical biology, 12(11), 908–910.

Becher, I. et al. (2018). Pervasive protein thermal stability variation during the cell cycle. Cell, 173(6),1495–1507.

Benjamini, Y. et al. (1995). Controlling the false discovery rate: a practical and powerful approach to multiple testing. Journal of the Royal statistical society: series B (Methodological), 57(1),289–300.

Berger, J. O. et al. (2005). Posterior model probabilities via path-based pairwise priors. Statistica Neerlandica, 59(1),3–15.

Berger, J. O. et al. (2001). Objective bayesian analysis of spatially correlated data. Journal of the American Statistical Association, 96(456),1361–1374.

Berger, J. O. et al. (2014). A bayesian approach to subgroup identification. Journal of biopharmaceutical statistics, 24(1),110–129.

Betancourt, M. (2017). A conceptual introduction to hamiltonian monte carlo. arXiv preprint arXiv:170l.02434.

Boisvert, F.-M. et al. (2007). The multifunctional nucleolus. Nature reviews Molecular cell biology, 8(7), 574–585.

Boukouvalas, A. et al. (2018). Bgp: identifying gene-specific branching dynamics from singlecell data with a branching gaussian process. Genome biology, 19(1), 65.

Box, G. E. (1980). Sampling and bayes’ inference in scientific modelling and robustness. Journal of the Royal Statistical Society: Series A (General), 143(4), 383–404.

Bruno, S. et al. (1992). Different effects of staurosporine, an inhibitor of protein kinases, on the cell cycle and chromatin structure of normal and leukemic lymphocytes. Cancer research, 52(2),470–473.

Bürkner, P.-C. et al. (2017). brms: An r package for bayesian multilevel models using stan. Journal of statistical software, 80(1),1–28.

Cai, Y. et al. (2003). Identification of new subunits of the multiprotein mammalian trrap/tip60-containing histone acetyltransferase complex. Journal of Biological Chemistry, 278(44),42733–42736.

Carlin, B. P. et al. (1995). Bayesian model choice via markov chain monte carlo methods. Journal of the Royal Statistical Society: Series B (Methodological), 57(3),473–484.

Carpenter, B. et al. (2017). Stan: A probabilistic programming language. Journal of statistical software, 76(1).

Caslini, C. et al. (2019). Hdac7 regulates histone 3 lysine 27 acetylation and transcriptional activity at super-enhancer-associated genes in breast cancer stem cells. Oncogene, 38(39),6599–6614.

Chae, H.-J. et al. (2000). Molecular mechanism of staurosporine-induced apoptosis in osteoblasts. Pharmacological Research, 42(4), 373–381.

Chaloner, K. et al. (1995). Bayesian experimental design: A review. Statistical Science,pages 273–304.

Chan-Penebre, E. et al. (2015). A selective inhibitor of prmt5 with in vivo and in vitro potency in mcl models. Nature chemical biology, 11(6),432.

Chang, S. et al. (2020). Comparison of bayesian and frequentist multiplicity correction for testing mutually exclusive hypotheses under data dependence. Bayesian Analysis.

Childs, D. et al. (2019). Nonparametric analysis of thermal proteome profiles reveals novel drug-binding proteins. Molecular& Cellular Proteomics, 18(12), 2506–2515.

Chopin, N. et al. (2010). Properties of nested sampling. Biometrika, 97(3),741–755.

Choudhary, C. et al. (2009). Lysine acetylation targets protein complexes and co-regulates major cellular functions. Science, 325(5942),834–840.

Cloutier, P. et al. (2017). R2tp/prefoldin-like component ruvbll/ruvbl2 directly interacts with znhit2 to regulate assembly of u5 small nuclear ribonucleoprotein. Nature communications, 8(1), 1–14.

Cooke, E. J. et al. (2011). Bayesian hierarchical clustering for microarray time series data with replicates and outlier measurements. BMC bioinformatics, 12(1), 399.

Crook, O. et al. (2020). A semi-supervised bayesian approach for simultaneous protein sub-cellular localisation assignment and novelty detection. bioRxiv.

Crook, O. M. et al. (2019). Semi-supervised non-parametric bayesian modelling of spatial proteomics, arXiv preprint arXiv:1903.02909.

Crowder, M. J. et al. (1990). Analysis of repeated measures, volume 41. CRC Press.

De Oliveira, V. (2007). Objective bayesian analysis of spatial data with measurement error. Canadian Journal of Statistics, 35(2), 283–301.

Del?Angelica, E. C. et al. (1999). Ap-4, a novel protein complex related to clathrin adaptors. Journal of biological chemistry, 274(11),7278–7285.

Doyon, Y. et al. (2004). Structural and functional conservation of the nua4 histone acetyl-transferase complex from yeast to humans. Molecular and cellular biology, 24(5),1884–1896.

Duane, S. et al. (1987). Hybrid monte carlo. Physics letters B, 195(2),216–222.

Dudley, R. et al. (1973). Sample functions of the gaussian process. The Annals of Probability, 1(1), 66–103.

Dziekan, J. M. et al. (2019). Identifying purine nucleoside phosphorylase as the target of quinine using cellular thermal shift assay. Science translational medicine, 11(473), eaau3174.

Dziekan, J. M. et al. (2020). Cellular thermal shift assay for the identification of drug–target interactions in the plasmodium falciparum proteome. Nature Protocols, pages 1–41.

Efron, B. (2004). Large-scale simultaneous hypothesis testing: the choice of a null hypothesis. Journal of the American Statistical Association, 99(465), 96–104.

Efron, B. (2012). Large-scale inference: empirical Bayes methods for estimation, testing, and prediction, volume 1. Cambridge University Press.

Feng, Y. et al. (2014). Global analysis of protein structural changes in complex proteomes. Nature biotechnology, 32(10), 1036.

Franken, H. et al. (2015). Thermal proteome profiling for unbiased identification of direct and indirect drug targets using multiplexed quantitative mass spectrometry. Nature protocols, 10(10), 1567.

Frottin, F. et al. (2019). The nucleolus functions as a phase-separated protein quality control compartment. Science, 365(6451), 342–347.

Fuglstad, G.-A. et al. (2019). Constructing priors that penalize the complexity of gaussian random fields. Journal of the American Statistical Association, 114(525), 445–452.

Gad, H. et al. (2014). Mth1 inhibition eradicates cancer by preventing sanitation of the dntp pool. Nature, 508(7495),215–221.

Geladaki, A. et al. (2019). Combining lopit with differential ultracentrifugation for high-resolution spatial proteomics. Nature communications, 10(1), 1–15.

Gelfand, A. E. et al. (1994). Bayesian model choice: asymptotics and exact calculations. Journal of the Royal Statistical Society: Series B (Methodological), 56(3),501–514.

Gelman, A. et al. (2006). Prior distributions for variance parameters in hierarchical models (comment on article by browne and draper). Bayesian analysis, 1(3), 515–534.

Gelman, A. et al. (1998). Simulating normalizing constants: From importance sampling to bridge sampling to path sampling. Statistical science, pages 163–185.

Gelman, A. et al. (2013). Bayesian data analysis. CRC press.

Gelman, A. et al. (2019). R-squared for bayesian regression models. The American Statistician, 73(3),307–309.

Ghosh, J. K. et al. (2003). Bayesian nonparametrics. Springer Science& Business Media.

Glasbey, C. (1979). Correlated residuals in non-linear regression applied to growth data. Journal of the Royal Statistical Society: Series C (Applied Statistics), 28(3),251–259.

Glasbey, C. (1980). Nonlinear regression with autoregressive time series errors. Biometrics,pages 135–139.

Grozinger, C. M. et al. (1999). Three proteins define a class of human histone deacetylases related to yeast hda1p. Proceedings of the National Academy of Sciences, 96(9),4868–4873.

Hirst, J. et al. (1999). Characterization of a fourth adaptor-related protein complex. Molecular biology of the cell, 10(8),2787–2802.

Hoffman, M. D. et al. (2014). The no-u-turn sampler: adaptively setting path lengths in hamiltonian monte carlo. Journal of Machine Learning Research, 15(1), 1593–1623.

Holmes, S. et al. (2018). Modern statistics for modern biology. Cambridge University Press.

Huang, J. X. et al. (2019). High throughput discovery of functional protein modifications by hotspot thermal profiling. Nature methods, 16(9),894–901.

Hubbert, C. et al. (2002). Hdac6 is a microtubule-associated deacetylase. Nature, 417(6887), 455–458.

Huber, K. V. et al. (2015). Proteome-wide drug and metabolite interaction mapping by thermal-stability profiling. Nature methods, 12(11), 1055–1057.

Ikura, T. et al. (2000). Involvement of the tip60 histone acetylase complex in dna repair and apoptosis. Cell, 102(4), 463–473.

Izaurralde, E. et al. (1994). A nuclear cap binding protein complex involved in pre-mrna splicing. Cell, 78(4), 657–668.

Izaurralde, E. et al. (1995). A cap-binding protein complex mediating u snrna export. Nature, 376(6542), 709–712.

Jafari, R. et al. (2014). The cellular thermal shift assay for evaluating drug target interactions in cells. Nature protocols, 9(9),2100.

Jarzab, A. et al. (2020). Meltome atlas—thermal proteome stability across the tree of life. Nature methods, 17(5), 495–503.

Johnson, R. et al. (2015). Sysbions: nested sampling for systems biology. Bioinformatics, 31(4), 604–605.

Justice, S. A. P. et al. (2020). Mutant thermal proteome profiling for characterization of missense protein variants and their associated phenotypes within the proteome. Journal of Biological Chemistry, pages jbc–RA120.

Karaman, M. W. et al. (2008). A quantitative analysis of kinase inhibitor selectivity. Nature biotechnology, 26(1),127–132.

Kass, R. E. et al. (1995). Bayes factors. Journal of the american statistical association, 90(430), 773–795.

Kirk, P. et al. (2012). Bayesian correlated clustering to integrate multiple datasets. Bioinformatics, 28(24), 3290–3297.

Kirk, P. D. et al. (2009). Gaussian process regression bootstrapping: exploring the effects of uncertainty in time course data. Bioinformatics, 25(10), 1300–1306.

Laubach, J. P. et al. (2015). Panobinostat for the treatment of multiple myeloma. Clinical cancer research, 21(21),4767–4773.

Leuenberger, P. et al. (2017). Cell-wide analysis of protein thermal unfolding reveals determinants of thermostability. Science, 355(6327), eaai7825.

Lewis, S. M. et al. (1997). Estimating bayes factors via posterior simulation with the laplace—metropolis estimator. Journal of the American Statistical Association, 92(438), 648–655.

Li, Y. et al. (2016). Hdacs and hdac inhibitors in cancer development and therapy. Cold Spring Harbor perspectives in medicine, 6(10), a026831.

Määttä, T. A. et al. (2020). Aggregation and disaggregation features of the human proteome. Molecular systems biology, 16(10), e9500.

Mateus, A. et al. (2016). Thermal proteome profiling: unbiased assessment of protein state through heat-induced stability changes. Proteome science, 15(1), 13.

Mateus, A. et al. (2018). Thermal proteome profiling in bacteria: probing protein state in vivo. Molecular systems biology, 14(7).

Mateus, A. et al. (2020). Thermal proteome profiling for interrogating protein interactions. Molecular Systems Biology, 16(3), e9232.

Meng, X.-L. et al. (2002). Warp bridge sampling. Journal of Computational and Graphical Statistics, 11(3), 552–586.

Meng, X.-L. et al. (1996). Simulating ratios of normalizing constants via a simple identity: a theoretical exploration. Statistica Sinica, pages 831–860.

Molina, D. M. et al. (2013). Monitoring drug target engagement in cells and tissues using the cellular thermal shift assay. Science, 341(6141),84–87.

Morohashi, Y. et al. (2010). Phosphorylation and membrane dissociation of the arf exchange factor gbfl in mitosis. Biochemical Journal, 427(3),401–412.

Mulvey, C. M. et al. (2017). Using hyperlopit to perform high-resolution mapping of the spatial proteome. Nature protocols, 12(6), 1110–1135.

Oates, M. E. et al. (2012). D2p2: database of disordered protein predictions. Nucleic acids research, 41(D1), D508–D516.

Obri, A. et al. (2014). Anp32e is a histone chaperone that removes h2a. z from chromatin. Nature, 505(7485), 648–653.

Ogawa, Y. et al. (2010). Development of a novel selective inhibitor of the down syndrome-related kinase dyrk1a. Nature communications, 1, 86.

Papadopoulos, C. et al. (2011). Splice variants of the dual specificity tyrosine phosphorylation-regulated kinase 4 (dyrk4) differ in their subcellular localization and catalytic activity. Journal of Biological Chemistry, 286(7),5494–5505.

Paulo, R. et al. (2005). Default priors for gaussian processes. The Annals of Statistics, 33(2), 556–582.

Perrin, J. et al. (2020). Identifying drug targets in tissues and whole blood with thermal-shift profiling. Nature Biotechnology, 38(3),303–308.

Piazza, I. et al. (2018). A map of protein-metabolite interactions reveals principles of chemical communication. Cell, 172(1-2),358–372.

Potel, C. M. et al. (2020). Impact of phosphorylation on thermal stability of proteins. bioRxiv.

Powell, J. L. (1994). Estimation of semiparametric models. Handbook of econometrics, 4, 2443–2521.

Queiroz, R. M. et al. (2019). Comprehensive identification of rna–protein interactions in any organism using orthogonal organic phase separation (oops). Nature biotechnology, 37(2), 169–178.

Ramsay, J. O.(2004). Functional data analysis. Encyclopedia of Statistical Sciences, 4.

Ramsay, J. O. et al. (1991). Some tools for functional data analysis. Journal of the Royal Statistical Society: Series B (Methodological), 53(3),539–561.

Rasmussen, C. E. (2003). Gaussian processes in machine learning. In Summer School on Machine Learning, pages 63–71. Springer.

Reid, J. E. et al. (2016). Pseudotime estimation: deconfounding single cell time series. Bioinformatics, 32(19),2973–2980.

Reinhard, F. B. et al. (2015). Thermal proteome profiling monitors ligand interactions with cellular membrane proteins. Nature methods, 12(12),1129–1131.

Robert, C. P. et al. (2009).Computational methods for bayesian model choice. In Aip conference proceedings, volume 1193, pages 251–262. American Institute of Physics.

Rümenapp, U. et al. (1997). Characteristics of protein-kinase-c-and adp-ribosylation-factor-stimulated phospholipase d activities in human embryonic kidney cells. European journal of biochemistry, 248(2),407–414.

Saei, A. A. et al. (2018). System-wide identification of enzyme substrates by thermal analysis (siesta). bioRxiv, page 423418.

Savitski, M. M. et al. (2014). Tracking cancer drugs in living cells by thermal profiling of the proteome. Science, 346(6205),1255784.

Savitski, M. M. et al. (2018). Multiplexed proteome dynamics profiling reveals mechanisms controlling protein homeostasis. Cell, 173(1), 260–274.

Schellman, J. A. (1994). The thermodynamics of solvent exchange. Biopolymers: Original Research on Biomolecules, 34(8),1015–1026.

Schopper, S. et al. (2017). Measuring protein structural changes on a proteome-wide scale using limited proteolysis-coupled mass spectrometry, nature protocols, 12(11), 2391.

Scott, J. G. et al. (2006). An exploration of aspects of bayesian multiple testing. Journal of statistical planning and inference, 136(7),2144–2162.

Scott, J. G. et al. (2010). Bayes and empirical-bayes multiplicity adjustment in the variable-selection problem. The Annals of Statistics, pages 2587–2619.

Seto, E. et al. (2014). Erasers of histone acetylation: the histone deacetylase enzymes. Cold Spring Harbor perspectives in biology, 6(4), a018713.

Shin, J. J. et al. (2019). Determining the content of vesicles captured by golgin tethers using lopit-dc. bioRxiv, page 841965.

Shindoh, N. et al. (1996). Cloning of a human homolog of thedrosophila minibrain/rat dyrk gene from “the down syndrome critical region” of chromosome 21. Biochemical and biophysical research communications, 225(1),92–99.

Skilling, J. et al. (2006). Nested sampling for general bayesian computation. Bayesian analysis, 1(4), 833–859.

Smith, I. R. et al. (2020). Identification of phosphosites that alter protein thermal stability. bioRxiv.

Solin, A. et al. (2020). Hilbert space methods for reduced-rank gaussian process regression. Statistics and Computing, 30(2),419–446.

Soundararajan, M. et al. (2013). Structures of down syndrome kinases, dyrks, reveal mechanisms of kinase activation and substrate recognition.Structure, 21(6), 986–996.

Sridharan, S. et al. (2019). Proteome-wide solubility and thermal stability profiling reveals distinct regulatory roles for atp. Nature communications, 10(1),1–13.

Stegle, O. et al. (2010). A robust bayesian two-sample test for detecting intervals of differential gene expression in microarray time series. Journal of Computational Biology, 17(3),355–367.

Stein, M. L. (2012). Interpolation of spatial data: some theory for kriging. Springer Science& Business Media.

Strauss, M. E. et al. (2020). Gpseudoclust: deconvolution of shared pseudo-profiles at single-cell resolution. Bioinformatics, 36(5),1484–1491.

Tan, C. S. H. et al. (2018). Thermal proximity coaggregation for system-wide profiling of protein complex dynamics in cells. Science, 359(6380), 1170–1177.

Thul, P. J. et al. (2017). A subcellular map of the human proteome. Science, 356(6340).

van der Vaart, A. W. et al. (2009). Adaptive bayesian estimation using a gaussian random field with inverse gamma bandwidth. The Annals of Statistics, 37(5B), 2655–2675.

Vehtari, A. et al. (2017). Practical bayesian model evaluation using leave-one-out cross-validation and waic. Statistics and computing, 27(5),1413–1432.

Wang, J.-L. et al. (2016). Functional data analysis. Annual Review of Statistics and Its Application, 3, 257–295.

Werner, T. et al. (2012). High-resolution enabled tmt 8-plexing. Analytical chemistry, 84(16), 7188–7194.

Wood, M. A. et al. (2000). An atpase/helicase complex is an essential cofactor for oncogenic transformation by c-myc. Molecular cell, 5(2), 321–330.

Yu, X. et al. (2012). A structure-based mechanism for arfl-dependent recruitment of coatomer to membranes. Cell, 148(3),530–542.

Zerbino, D. R. et al. (2018). Ensembl 2018. Nucleic acids research, 46(D1),D754–D761.

